# A mass cytometry approach to track the evolution of T cell responses during infection and immunotherapy by paired T cell receptor repertoire and T cell differentiation state analysis

**DOI:** 10.1101/2024.01.11.575237

**Authors:** Jesse Garcia Castillo, Rachel DeBarge, Abigail Mende, Iliana Tenvooren, Diana M. Marquez, Adrian Straub, Dirk H. Busch, Matthew H. Spitzer, Michel DuPage

**Author notes:** These authors contributed equally (Jesse Garcia Castillo, Rachel DeBarge). These authors contributed equally (Matthew H. Spitzer, Michel DuPage). Contact info.

## Abstract

T cell receptor (TCR) recognition followed by clonal expansion is a fundamental feature of adaptive immune responses. Here, we developed a mass cytometric (CyTOF) approach combining antibodies specific for different TCR Vα– and Vβ-chains with antibodies against T cell activation and differentiation proteins to identify antigen-specific expansions of T cell subsets and assess aspects of cellular function. This strategy allowed for the identification of expansions of specific Vβ and Vα chain expressing CD8^+^ and CD4^+^ T cells with varying differentiation states in response to *Listeria monocytogenes*, tumors, and respiratory influenza infection. Expanded Vβ chain expressing T cells could be directly linked to the recognition of specific antigens from *Listeria*, tumor cells, or influenza. In the setting of influenza infection, we showed that the common therapeutic approaches of intramuscular vaccination or convalescent serum transfer altered the clonal diversity and differentiation state of responding T cells. Thus, we present a new method to monitor broad changes in TCR specificity paired with T cell differentiation during adaptive immune responses.

## Introduction

The basis of adaptive immunity is the clonal expansion of rare T or B cells that are specifically reactive to unique antigens. In response to an infection, CD4^+^ or CD8^+^ T cells possessing a T cell receptor (TCR) that recognizes an antigen from the pathogen can activate, proliferate, differentiate, and then migrate to the site of infection to control it^1–3^. Clonal expansions include 3-10 cycles of division, creating 10 to 1,000+ cells from a single reactive T cell^4–7^. During the proliferative phase, and depending on the type of insult, clonally-expanded T cells will differentiate, gaining distinct functions with some acquiring the capacity to form long-lived memory cells^8–11^. While immune-based therapies can change the function of T cells to better fight disease, it has also been shown that they can change the clonal diversity (or repertoire) of the responding T cells^12–17^.

Although methods to track clonal T cell responses exist, current strategies are either slow and labor intensive or lack broad applicability. Bulk TCR sequencing can identify patterns in the use of T cell receptor genes but cannot directly link phenotypic or functional information about the cells that comprise clonal expansions of T cells^14,18,19^. Single cell sequencing can pair unique TCR sequences with gene expression analysis to infer phenotype and function; however, it is limited in the number of cells that can be analyzed^20–22^. Furthermore, transcription may infer some phenotypic attributes of T cells, but it is often imperfect, as many key cellular processes are regulated post-transcriptionally^23^. Functional assays, such as ELISPOT, can enumerate cytokine production from antigen-specific cells but are limited to measuring one function at a time^24^. Tetramer staining can identify T cells recognizing a particular peptide:MHC combination but require custom reagents for each specificity, which prevents investigation of the total reactive T cell pool^25,26^. Most challenging for these functional assays is the requirement that the antigens recognized by T cells already be known; thus, studying broad changes in the clonality and T cell responses together with measurements of T cell function has not been possible. Thus, we developed an approach to balance the trade-offs in T cell characterization techniques. We hypothesized that we could use simultaneous staining of unique Vα and Vβ chains on T cells to learn about the TCR repertoire during a response. While not tracking single T cell clones, this approach would be fast, iterative, and scalable for use in many disease settings in mice and humans.

In this study, we used a novel mass cytometry by time of flight (CyTOF) approach to resolve these challenges. CyTOF is a technology that enables the quantification of >45 proteins in millions of single cells^27^. The increase in the number of proteins that can be analyzed at once allowed us to develop antibody panels to detect both the Vα and Vβ chains of different TCRs to monitor changes in the TCR repertoire while simultaneously measuring proteins to delineate T cell differentiation and function^28^. Flow cytometry based staining of Vα and Vβ TCR chains has been done previously to track clonal populations, but limitations of fluorescence-based readouts required the use of multiple panels or sacrificed the amount of phenotypic information gathered simultaneously^29–31^. As a demonstration of the utility of our approach, we linked expansions in TCR Vα and Vβ chains to clonal expansions of T cells responding to well-defined antigens from *Listeria monocytogenes* (*Lm*) and influenza. While our approach does not require that the antigens targeted by T cells be known to track changes in the TCR repertoire, these studies allowed us to benchmark its performance in contexts of defined immunodominant antigens while also examining additional unknown T cell responses. Using our strategy to study different treatment modalities for respiratory infection by influenza, we show that prophylactic vaccination by intramuscular injection versus passive transfer of convalescent serum both led to changes in the repertoire of responding T cells as well as in the differentiation state of T cells utilizing specific TCR Vα and Vβ chains. By testing our approach in the setting of influenza infection, *Lm* infection, and cancer, we validate a new strategy that is readily applicable to study the T cell response in multiple settings without the need for custom reagents or tailored biological systems. We also developed automated data analysis approaches to identify populations of T cells with particular TCR Vα and Vβ chains. Our findings advance the understanding of T cell priming and fate determination, elucidate ways immunotherapies could be improved, and enable future studies through technological development and validation.

## Results

### Tracking the phenotypes of clonal T cell responses against *Listeria monocytogenes* with CyTOF

In this study, we aimed to test whether mass cytometry (CyTOF) could be used to track clonal expansions of T cells during an immune response by using antibodies against TCR Vβ and Vα chains combined with highly-multiplexed measurements of the protein expression of phenotypic markers of T cell activation and differentiation. First, we injected mice with attenuated strains of *Listeria monocytogenes,* referred to as attenuated double-deleted (*LADD*, *ΔActAΔInlB*), that either expressed the common model antigen Ovalbumin (*LADD-OVA*) or did not (*LADD*) compared to PBS-injected controls (Figure 1a)^32^. We chose *LADD* because it has been widely used to safely generate robust T cell responses, especially CD8^+^ T cell responses, against overexpressed antigens in *LADD* as a cancer vaccine strategy^32,33^. Indeed, five days after administration, we observed increased activation of CD8^+^ and CD4^+^ T cells in the spleens of *LADD*-infected mice compared to controls (Extended Data Figures 1a,1b and 2a). Clustering analysis of CD8^+^ T cells based on activation and differentiation proteins visualized by Uniform Manifold Approximation and Projection (UMAP) revealed effector T cell (T_eff_) and some effector memory T cell (T_em_) populations unique to *LADD*-infected mice (Figure 1b, Extended Data Table 2). Due to the lack of CD62L in our initial panels, we were not able to distinguish T_cm_ and T_nv_ populations during *Listeria* infection. The T_eff_ populations were defined by the expression of LFA-1/CD11a and T-bet (Figure 1c). However, the Teff_1 population stood out as particularly different due to high expression of Ki-67 and PD-1. This cluster also expressed proteins CD49b, ICOS, and CD25 (Figures 1c and 1d), suggesting it consisted of early activated, antigen-specific CD8^+^ T cells (T_ea_) that we recently described to have a unique metabolic state and to peak in abundance at this time after *Listeria* infection^34^. Similar observations were made in CD4^+^ T cells, where Ki-67 expression differentiated two T_eff_ populations (Extended Data Figures 2a-2c). Given that the CD8^+^ Teff_1 population had the greatest enrichment of markers indicative of antigen experience, we utilized Ki-67 and PD-1 expression to look for enrichment of TCR Vβ and Vα chains in *LADD-* and *LADD-OVA*-infected mice within this proliferating sub-population (Figure 1e). We identified Vβ5.1/2^+^ and Vβ12^+^ T cell populations expanded in mice infected with *LADD* or *LADD-OVA*, whereas Vβ14^+^ T cells were only expanded in mice infected with *LADD-OVA* (Figure 1e-1h). Furthermore, only *LADD-OVA*-infected mice had an expansion of Vα2^+^ CD8^+^ T cells that were also Vβ14^+^ (Figure 1e, 1i). The expansion of Vβ14^+^Vα2^+^ T cells in *LADD-OVA*-infected mice may indicate that T cells with these TCR chains have clonally expanded due to their specificity for the OVA antigen. UMAP visualization and quantification of Vβ14 expression in CD8^+^ T cells, and more specifically Vβ14^+^Vα2^+^ CD8^+^ T cells, revealed a clear enrichment in T cells expressing these TCR chains in the Teff_1 cluster only after *LADD-OVA* infection (Figure 1j, 1k, Extended Data Figure 1c), whereas Vβ12^+^ CD8^+^ T cells could be found expanded in both *LADD-* and *LADD-OVA*-infected mice (Extended Data Figure 1d). In conventional CD4^+^ T cells, we also observed increased usage of specific TCR Vβ and Vα chains in mice infected with *LADD* or *LADD-OVA*; however, no expansions were unique to *LADD-OVA* (Extended Data Figure 2e-2h). In particular, we found that Vβ13^+^Vα2^+^ CD4^+^ T cells were dramatically increased in both *LADD*-infected mice (Extended Data Figure 2i). Again, UMAP visualization of TCR Vβ and Vα chains revealed a clear enrichment in CD4^+^ T cells expressing either Vβ13 and Vα2, or Vβ14 in the Teff_1 cluster of CD4^+^ T cells with *LADD-*infection (Extended Data Figures 2j-2m). These results show that changes in the frequency of TCR Vβ and Vα chains used by T cells can be detected and quantified by our CyTOF approach during an immune response to *LADD*. We thus hypothesized that these changes in TCR chain usage were driven by clonal T cell expansions of CD8^+^ and CD4^+^ T cells directed against specific antigens expressed by *LADD*.

**Figure 1.**
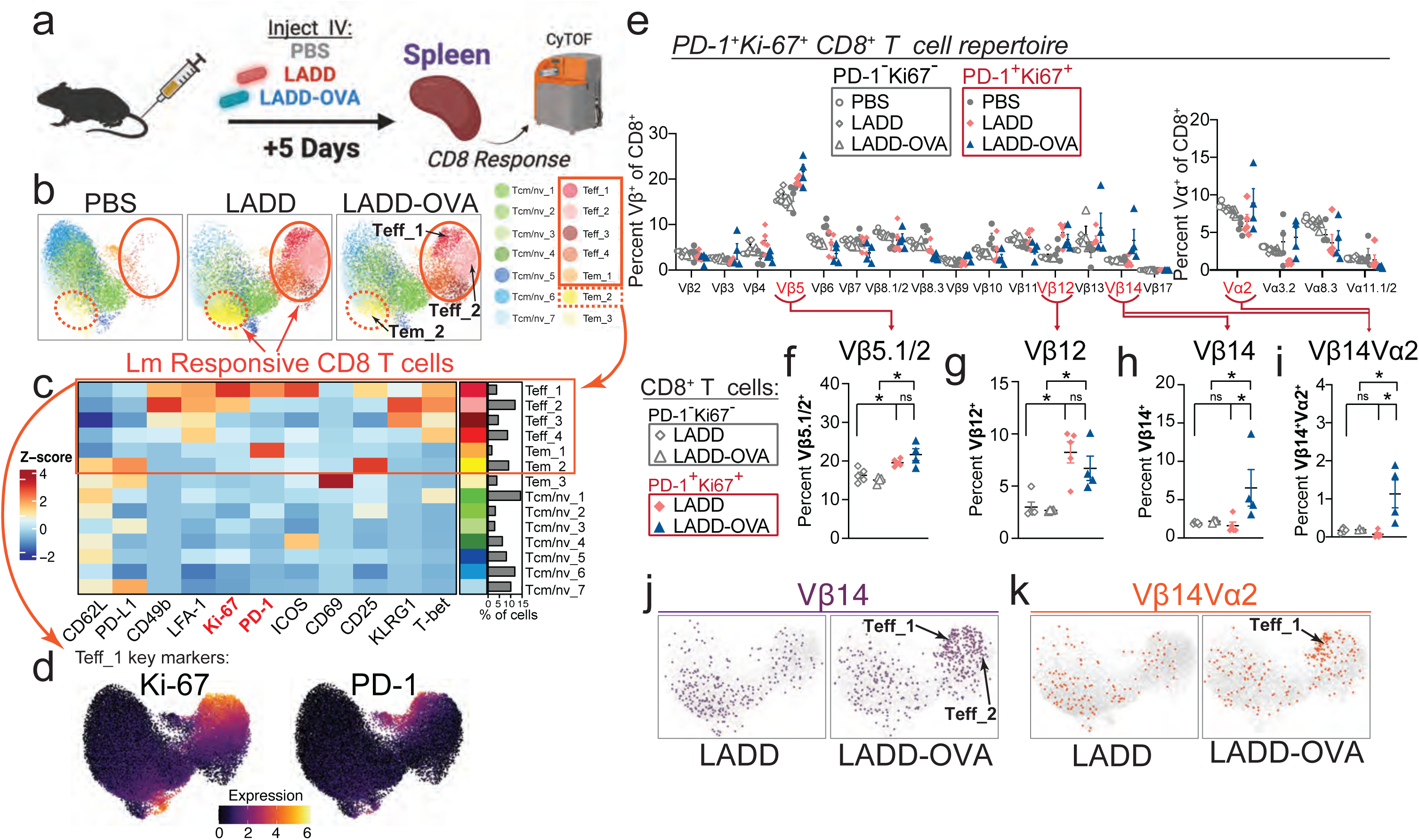
| Tracking the phenotypes of clonal T cell responses against attenuated *Listeria (LADD)* with CyTOF. (a) C57BL/6J mice were injected with *LADD, LADD-OVA, or PBS* intravenously (i.v.) and after 5 days splenocytes were analyzed by CyTOF. (b) UMAP visualization of CD8^+^ T cell clusters based on expression of non-TCR proteins. (c) Heatmap of non-TCR protein expression annotated by cluster and the percentage of cells falling into each cluster. (d) UMAP visualization of CD8^+^ T cells colored by the expression of PD-1 and Ki-67 in *LADD*-infected mice. (e) Frequency of PD-1^+^Ki-67^+^ and PD-1^−^Ki-67^−^ CD8^+^ T cells using specific TCR Vβ (left) or Vα chains (right) in response to PBS, *LADD*, or *LADD-OVA*. (f-i) Frequency of Vβ5.1/2^+^ (f), Vβ12^+^ (g), Vβ14^+^ (h), or Vβ14^+^Vα2^+^ among PD-1^+^Ki-67^+^ versus PD-1^−^Ki-67^−^ CD8^+^ T cells in *LADD* or *LADD-OVA* infected mice. (j-k) UMAP visualization of pooled CD8^+^ T cells colored by the expression of Vβ14^+^ (j) and Vβ14^+^Vα2^+^ (k) in *LADD* or *LADD-OVA* infected mice. Results from n=5 for PBS, n=4 for *LADD*, n=4 for *LADD-OVA*, *P<0.05, **P<0.01, ***P<0.001 by unpaired two-tailed Student’s t-test, mean ± s.e.m.

### Expanded CD8^+^ and CD4^+^ T cells expressing specific TCR Vβ chains are enriched for recognition of antigens from *LADD*

To validate that the increased CD8^+^ and CD4^+^ T cells with specific TCR Vβ and Vα chains was the result of their recognition of antigens expressed by *LADD* (OVA or *Listeria-*derived antigens), we tested known antigens in peptide restimulation assays and loaded MHC tetramers with defined peptides to stain antigen-specific T cells (Figure 2a and Extended Data Figures 3a-3c and 4a,4b). We calculated a fold-enrichment for either interferon-gamma (IFN-γ)-producing (Figure 2b, 2d, 2f, and 2h) or tetramer-positive (Figure 2c and 2g) T cells for each specific TCR Vβ chain as compared to all CD8^+^ or CD4^+^ T cells. The major epitope from OVA recognized by H-2K^b^-restricted CD8^+^ T cells is the peptide SIINFEKL^35^. From our analysis in Figure 1, we hypothesized that the expanded Vβ14^+^ CD8^+^ T cells were specific for OVA and most likely the SIINFEKL epitope. In support of our hypothesis, SIINFEKL peptide stimulation of splenocytes from *LADD-OVA*-infected mice led to IFN-γ production in CD8^+^ T cells, and IFN-γ^+^ cells were significantly enriched for the expression of the Vβ14 TCR (∼10-fold) compared to all CD8^+^ T cells or CD8^+^ T cells expressing other TCR Vβ chains (Figure 2b and Extended Data Figure 3d,3e). CD8^+^ T cells using Vβ5.1/2 were also enriched in this experiment, although very weakly. Notably, Vβ5 is the TCR chain used in the OT-1 TCR transgenic mouse that recognizes the SIINFEKL epitope presented on H-2K^b^ MHC complexes^36^. Using SIINFEKL-loaded H-2K^b^ tetramers, we also found that SIINFEKL-tetramer-positive cells were significantly enriched for Vβ14 usage, and even further enriched in Vβ14^+^Vα2^+^ cells (Figure 2c). These results indicate that Vβ14^+^ and Vβ14^+^Vα2^+^ CD8^+^ T cells in *LADD-OVA*-infected mice are significantly enriched in cells that recognize and respond to the SIINFEKL antigen, consistent with our hypothesis that we can use Vβ14-expression together with activation and proliferation markers as a surrogate for CD8^+^ T cells specific to OVA in *LADD-OVA* infected mice.

**Figure 2.**
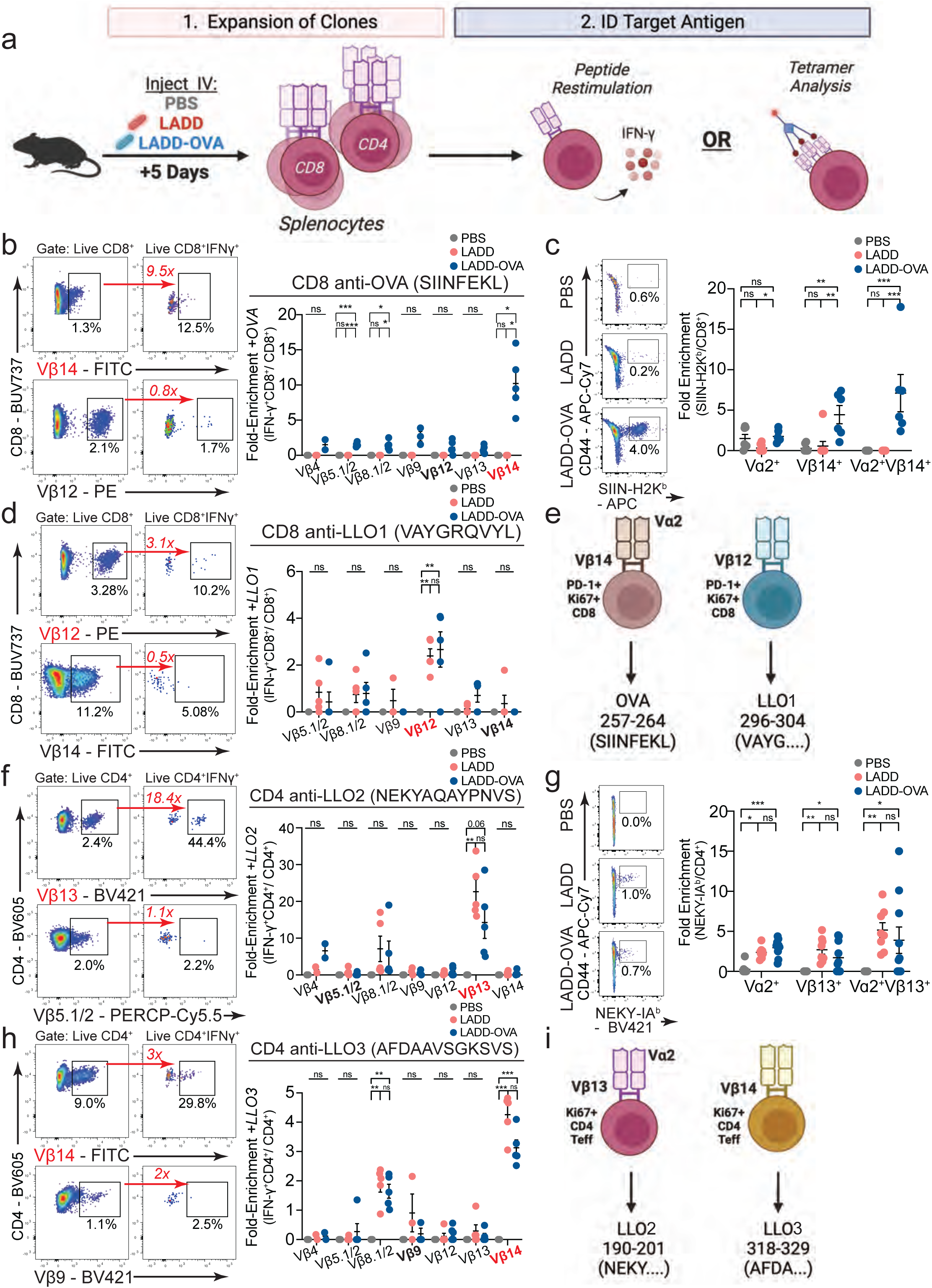
| The OVA– and *LADD*-expanded CD8^+^ and CD4^+^ T cells recognize *LADD*-encoded antigens. (a) C57BL/6J mice were injected with *LADD, LADD-OVA, or PBS* i.v., and 5 days later, antigen recognition was analyzed by peptide restimulation or tetramer staining. (b) Representative flow plots and quantification of fold-enrichment in Vβ-chain usage by IFNγ^+^CD8^+^/CD8^+^ T cells with OVA (SIINFEKL) peptide restimulation. (c) Representative flow plots and quantification for fold-enrichment using SIINFEKL-loaded H-2K^b^ tetramer staining. (d) Representative flow plots and quantification of fold-enrichment in Vβ-chain usage by IFNγ^+^CD8^+^/CD8^+^ T cells with LLO1 (VAYGRQVYL) peptide restimulation. (e) Schematic summarizing CD8^+^ T cells using specific Vβ-chains enriched to recognize specific antigens from LADD. (f) Representative flow plots and quantification of fold-enrichment in Vβ-chain usage by IFNγ^+^CD4^+^/CD4^+^cells with LLO2 (NEKYAQAYPNVS) peptide restimulation. (g) Representative flow plots and quantification for fold-enrichment in tetramer assay using IA^b^-NEKYAQAYPNVS tetramer staining. (h) Representative flow plots and quantification of fold-enrichment in Vβ-chain usage by IFNγ^+^CD4^+^/CD4^+^ T cells with LLO3 (AFDAAVSGKSVS) peptide restimulation. (i) Schematic summarizing CD4^+^ T cells using specific Vβ-chains enriched to recognize specific antigens from LADD. Results from n=2 independent experiments, *P<0.05, **P<0.01, ***P<0.001 by One-Way ANOVA for peptide restimulation assays and unpaired two-tailed Student’s t-test for tetramer assays, mean ± s.e.m.

Next, we tested whether the CD8^+^ and CD4^+^ T cells that expanded with specific TCR Vβ chains with both *LADD* and *LADD-OVA* infection recognized *Listeria*-derived antigens. Several *Listeria* antigens recognized by CD8^+^ and CD4^+^ T cells are derived from the Listeriolysin O (LLO) protein^37^. We tested one peptide from LLO that is presented on H-2K^b^ to CD8^+^ T cells (VAYGRQVYL, denoted LLO1) and two additional peptides presented on MHC-II (I-A^b^) to CD4^+^ T cells, which we called LLO2 (NEKYAQAYPNVS) and LLO3 (AFDAAVSGKSVS)^38^. Splenocytes from *LADD* or *LADD-OVA* injected mice were stimulated with VAYGRQVYL, enabling us to identify a population of CD8^+^ T cells that produced IFN-γ. Within the IFN-γ^+^ population, CD8^+^ T cells expressing Vβ12 were enriched ∼3-fold compared to all CD8^+^ T cells (Figure 2d and Extended Data Figures 2f,2g). Therefore, we identified Vβ14^+^Vα2^+^ CD8^+^ T cells recognizing SIINFEKL and Vβ12^+^ CD8^+^ T cells recognizing VAYGRQVYL (Figure 2e).

To investigate the CD4^+^ T cell response to *LADD*, splenocytes from *LADD*-infected mice were stimulated with LLO2. In these mice, CD4^+^Vβ13^+^ T cells were enriched ∼20-fold within the IFN-γ^+^ population compared to all CD4^+^ T cells or compared to other Vβ-expressing CD4^+^ T cells (Figure 2f and Extended Data Figures 4c,4d). Using MHC-II tetramers loaded with LLO2 (NEKY…), we also observed a ∼5-fold enrichment in Vβ13^+^Vα2^+^ CD4^+^ T cells recognizing LLO2 compared to all CD4^+^ T cells from *LADD*-infected mice (Figure 2g). Using the LLO3 peptide (AFDA…) to stimulate splenocytes from infected mice, we observed a ∼3-fold enrichment in Vβ14^+^ CD4^+^ T cells in the IFN-γ^+^ population compared to all CD4^+^ T cells or compared to other Vβ-expressing CD4^+^ T cells (Figure 2h and Extended Data Figures 4e,4f). Interestingly, we also noted a significant but more modest enrichment in Vβ8.1/2^+^ CD4^+^ T cells in the IFN-γ^+^ population, which was also weakly observed with LLO2 stimulation. Thus, in several cases, enrichment in proliferating T cells expressing a specific Vβ chain was sufficient to capture expansions in CD8^+^ and CD4^+^ T cells with TCRs specific for antigens expressed by *LADD* (Figures 2e and 2i). Therefore, measuring changes in TCR Vβ and Vα chain use within the population of proliferating cells can identify clonally expanded T cells during an immune response. Our approach expands the use of CyTOF to identify antigen-responsive T cells at a single cell level not possible by bulk T cell analysis, at a magnitude and throughput not attainable by single-cell TCR sequencing, and without requiring prior knowledge of the antigen(s) of interest.

To compare our CyTOF method to detect and phenotype antigen-responsive clonal populations to single-cell RNA sequencing methods, we re-analyzed a recently acquired single-cell RNA-seq dataset from CD8^+^ T cells sorted from spleens of mice infected with OVA-expressing Listeria^39^. Cells in this study were sorted that exhibited an activated (CD44^+^) phenotype or were SIINFEKL-reactive by tetramer staining. Phenotypic analysis of the CD8^+^ T cells identified six distinct clusters of cells, which expressed similar markers as compared to our results using our CyTOF method (Extended Data Figures 5a-5c). Expression of *Mki67* and *Pdcd1* was also concentrated in clusters that were annotated as T_eff_ and T_em_ lineages (Extended Figure 5d). Next, we determined whether Vβ14^+^ and Vβ14^+^Vα2^+^ CD8^+^ T cells were present among the SIINFEKL-reactive CD8^+^ T cells in this dataset. Indeed, CD8^+^ T cells expressing *Trbv31* (Vβ14) and *Trav14d-2* (Vα2.1) were enriched in both the T_eff_ and T_em_ clusters that were determined to be SIINFEKL-reactive (Extended Data Figures 5e-5f). When quantifying this enrichment, *Trbv31* (Vβ14) cells were enriched only in the SIINFEKL-reactive cells (Extended Data Figure 5g, left), but not in the cells that didn’t react to SIINFEKL (Extended Data Figure 5g, right). Most importantly, SIINFEKL-reactive CD8^+^ T cells that expressed *Trav14d-2* (Vα2.1) were dramatically enriched in *Trbv31* (Vβ14) expression (Extended Data Figure 5h). Therefore, our CyTOF method and scRNA-seq methods both captured the same enrichment in TCR chain usage and CD8^+^ T cell phenotypes in response to *Listeria* overexpressing OVA.

Next, we compared our results with this scRNA-seq dataset for their overall capacity to capture the distribution of TCR chain usage in this setting. First, we directly compared all Vβ chain assignments between both methods and observed comparable frequencies of Vβ chain assignment between the two methods (Extended Data Figure 5i-5j). When the two datasets were compared after removing unassigned TCRs in each dataset, the concordance was even higher for Vβ chain use in both datasets (Extended Data Fig 5i). In fact, a direct comparison of Vβ chain abundances showed high correlation between both methods (Extended Data Figure 5k). Taken together, the comparison of our CyTOF method to a paired scRNA-seq dataset generated in a separate lab but with the same *Listeria* infection model, shows the feasibility and reproducibility of our CyTOF method for identifying and characterizing clonal T cell responses during infection.

### Activated Vβ14^+^ CD8^+^ T cells expand in response to OVA-expressing tumors and recognize the OVA antigen

Next, we investigated whether OVA expression in cancer cells would also lead to an expansion of Vβ14^+^ CD8^+^ T cells specific for OVA as observed with *LADD-OVA* infection (Figure 1). We inoculated mice with B16F10 melanoma tumor cells that either did or did not express OVA, and we monitored the frequency of Vβ14^+^ cells in mice^40^. In mice with B16F10-OVA tumors, we observed an increase in the frequency of Vβ14^+^ CD8^+^ T cells within the proliferating (Ki-67^+^PD-1^+^) cells from tumors and tumor draining lymph nodes (tdLN) of mice, while this was not observed in B16F10 tumors lacking OVA expression (Figure 3a). Using tetramer staining, we validated that SIIN-H-2K^b^ ^+^ cells were enriched in Vβ14^+^ CD8^+^ T cells (Figures 3b and 3c). There was a ∼2-fold enrichment in B16F10-OVA tumors and a ∼3-fold enrichment in the tdLN. These results demonstrate that our approach can broadly identify OVA-specific CD8^+^ T cells based on their use of the TCR Vβ14 chain across two different settings of immune responses.

**Figure 3.**
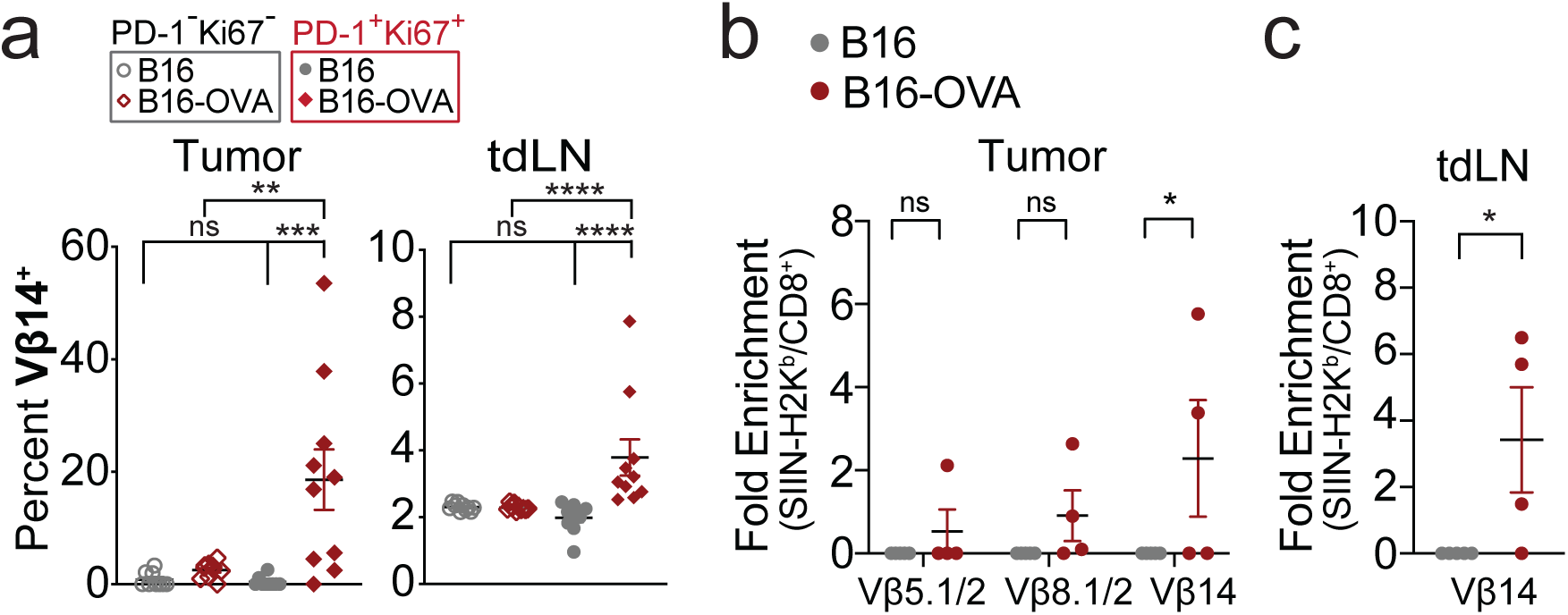
| Activated Vβ14^+^ CD8^+^ T cells expand in OVA-expressing tumor model. (a) Frequencies of Vβ14^+^ CD8^+^ T cells in B16F10 tumors (+/− OVA expression) and tumor-draining lymph nodes (tdLN) separated by non-proliferating (PD-1^−^Ki67^−^) vs. proliferating (PD-1^+^Ki67^+^) cells. (b) Quantification for fold-enrichment of H2-K^b^-SIINFEKL tetramer staining in tumors from B16F10 and B16F10-OVA tumor-bearing mice. (c) As in (b) except from tdLN. Results from n=10 for B16F10, n=10 for B16F10-OVA for CyTOF analysis and n=5 for B16F10, n=4 for B16F10-OVA for tetramer analysis, *P<0.05, **P<0.01, ***P<0.001 by unpaired two-tailed Student’s t-test for flow analysis in tumors and tdLNs and one-way ANOVA for tetramer analysis in tumors, mean ± s.e.m.

### Respiratory infection with influenza induces expansions of CD8^+^ and CD4^+^ T cells using specific Vβ chains linked to the recognition of influenza antigens

During localized infections at non-lymphoid sites, such as the lung during influenza infection, antigens from a pathogen are trafficked from the site of infection to secondary lymphoid structures to activate and expand T cells that express a TCR that recognizes the antigen. These expanded T cells then migrate and infiltrate pathogen-infected tissues and mount an appropriate inflammatory response to remove the pathogen^41^. To test whether our approach was capable of tracking T cell expansions in a natural model of respiratory infection, we used intranasal delivery of the PR8 strain of influenza (flu) and tracked expansions in TCR Vβ and Vα chain use in T cells. We performed our CyTOF analysis ten days after flu infection across multiple tissues: the lungs, mediastinal lymph node (medLN) that drains the lung tissue, non-draining inguinal lymph nodes (ndLN), and the spleen (Figure 4a). We observed increased activation of both CD8^+^ and CD4^+^ T cells by Ki-67 and PD-1 expression specifically in the lung, medLN, and spleen (Extended Data Figures 6a-6c). Clustering analysis of activation and differentiation proteins on CD8^+^ T cells visualized by UMAP showed the appearance of effector (T_eff_), effector memory (T_em_), and central memory (T_cm_) CD8^+^ T cell populations enriched in the flu-infected mice (Figure 4b, Extended Data Table 2). The two T_eff_ populations shared expression of key phenotypic markers (e.g., KLRG1 and T-bet) that defined their effector lineage, but had differential expression of functional markers (e.g., Granzyme B, IFN-γ). This differential expression of phenotypic and lineage markers has been previously shown for both the CD8 and CD4 compartment during influenza infection^42^. In comparison, T_em_ and T_cm_ subsets were defined by differential expression of canonical markers (e.g., CD62L, CD44, Slamf6). Similar observations were made in CD4^+^ T cells with the identification of T_eff_, T_em_, and T_cm_ subsets (Extended Data Figures 7a and 7b).

**Figure 4.**
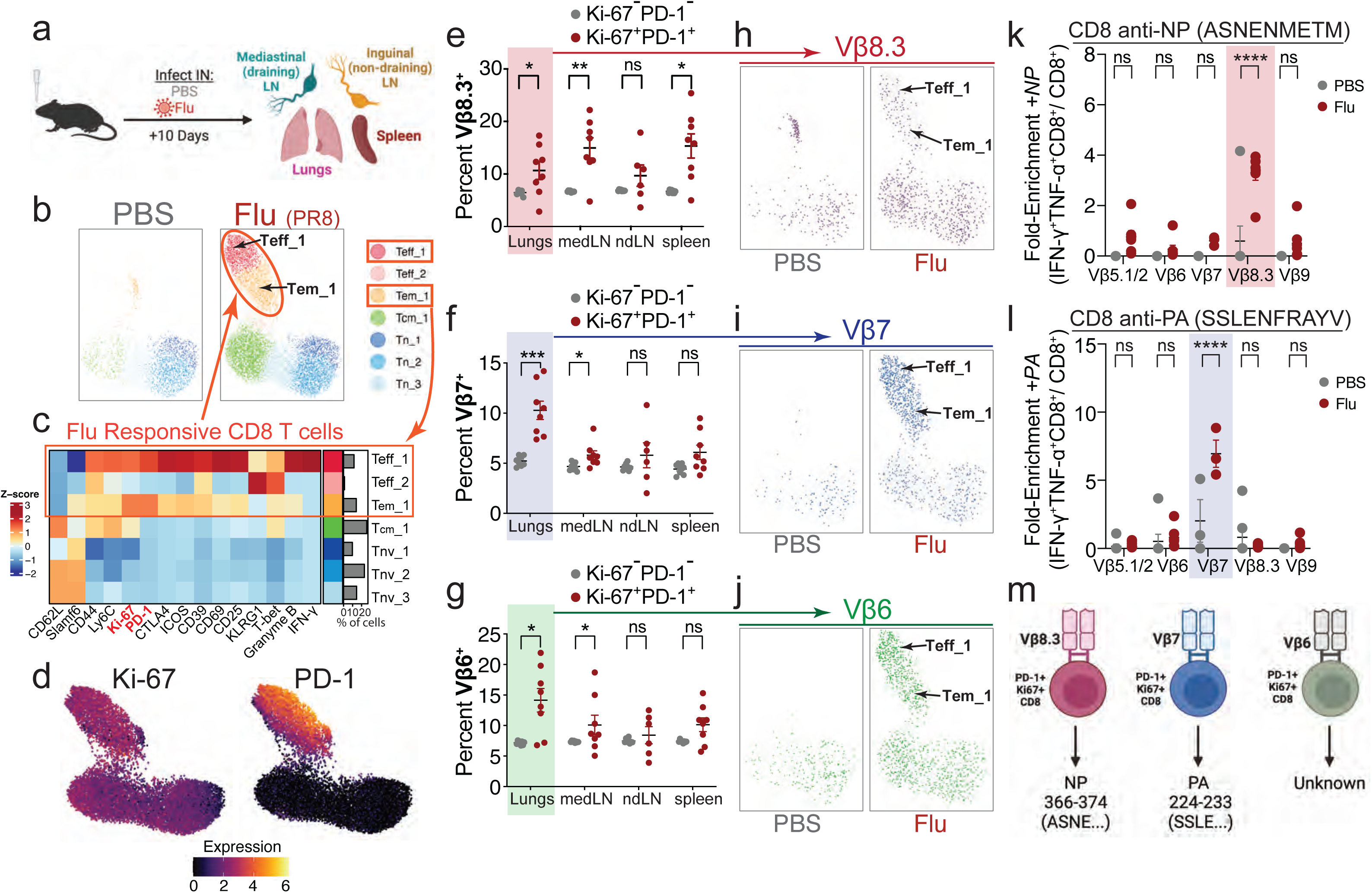
| Tracking the expansion and recognition of CD8^+^ T cells specific for influenza antigens across multiple tissues. (a) C57BL/6J mice were infected with *PR8 H1N1 influenza* (Flu) intranasally (IN), and indicated tissues were analyzed for T cell responses. (b) UMAP visualization of CD8^+^ T cell clusters based on expression of non-TCR proteins. (c) Heatmap of non-TCR protein expression annotated by cluster and fraction of cells falling into each cluster. (d) UMAP visualization of CD8^+^ T cells colored by the expression Ki-67 and PD-1 in flu-infected mice. (e-g) Comparison of the frequencies of CD8^+^ T cells using Vβ8.3 (e), Vβ7 (f), and Vβ6 (g) in non-proliferating (PD-1^−^Ki-67^−^) and proliferating (PD-1^+^Ki-67^+^) cells across tissues of flu-infected mice. (h-j) UMAP visualization of pooled CD8^+^ T cells colored by the expression of Vβ8.3 (h), Vβ7 (i), and Vβ6 (j) in PBS versus flu-infected mice. (k) Quantification of fold-enrichment in Vβ-chain usage by CD8^+^ T cells stimulated with the PR8 NP peptide (ASNENMETM). (l) Quantification of fold-enrichment in Vβ-chain usage by CD8^+^ T cells stimulated with the PR8 PA peptide (SSLENFRAYV). (m) Schematic summarizing CD8^+^ T cells using specific Vβ-chains enriched to recognize specific antigens from flu. Results from n=7 for PBS, n=7 for PR8, *P<0.05, **P<0.01, ***P<0.001 by unpaired two-tailed Student’s t-test for CyTOF analysis and one-way ANOVA for peptide restimulation assay, mean ± s.e.m.

Using a similar approach to *LADD* infection, we utilized Ki-67 and PD-1 expression to identify TCR Vβ and Vα chains associated with putative *flu*-specific CD8^+^ T cells (Figure 4d). We identified Vβ8.3, Vβ7, and Vβ6 TCR chains expanded in CD8^+^ T cells in the lungs of influenza infected mice (Figures 4e-4g). However, the expansion of these Vβ chains in CD8^+^ T cells was variable in the medLN, ndLN, and spleen of flu-infected mice. When we overlaid Vβ chain expression on UMAPs of CD8^+^ T cells, where dimensionality reduction was based only on expression of activation and differentiation markers but not Vβ or Vα chains, we found that Vβ8.3, Vβ7, and Vβ6 expression was enriched in the Teff_1 and Tem_1 compared to the T_cm_ and T_nv_ clusters (Figures 4h-4j). Next, we tested whether the expanded Vβ chain-expressing CD8^+^ T cells recognized any of the defined antigens from flu^43–45^. We prepared single cell suspensions of cells from the lungs of PBS or flu-infected mice and stimulated with the following peptides: ASNENMETM, derived from the PR8 nucleoprotein (NP) and SSLENFRAYV, derived from the viral polymerase subunit A (PA). There was ∼4-fold enrichment in Vβ8.3^+^ CD8^+^ T cells producing IFN-γ and TNF-α in response to ASNENMETM compared to all CD8^+^ T cells, even those expressing the other Vβ chains that expanded in flu-infected mice (Figure 4k). Single cells from lung tissue stimulated with SSLENFRAYV peptide exhibited ∼6-fold enrichment of Vβ7^+^ CD8^+^ T cells producing IFN-γ and TNF-α compared to all CD8^+^ T cells (Figure 4l).

For CD4^+^ T cells, there were two T_eff_ clusters that appeared in the lungs of infected mice (Extended Data Figure 7a). These clusters were distinguished by differential expression of Ly6C, Granzyme B, CD39, Slamf6, and CD69 (Extended Data Figure 7b). Using Ki-67 as a marker of proliferating cells, we observed the expansion of Vβ14^+^ CD4^+^ cells in the lungs, spleen, and in the medLN (Extended Data Figures 7c and 7d). The Vβ14^+^ CD4^+^ T cells that expanded were enriched in the Teff_1 cluster that had the highest expression of inflammatory and activation markers (Extended Data Figure 7e). We stimulated cells from PR8-or PBS-infected lungs with the LILRGSVAHKSCLPACV peptide, a NP-derived epitope presented on I-A^b^ and observed ∼5-fold enrichment in Vβ14^+^ CD4^+^ T cells compared to cells expressing other Vβ chains (Extended Data Figure 7f). Thus, Vβ14^+^ CD4^+^ T cells expanded in response to flu are enriched for recognition of the NP peptide (LILR…) (Extended Data Figure 7g).

Regulatory T cells (Tregs) have also been shown to impact outcomes of infection with flu^46^. Therefore, we tested whether our method allowed us to observe expansions in clonal Treg populations in response to flu infection. Interestingly, when looking across all tissues, we only observed an increase in the activation of Tregs in the medLN, but not in the lungs, ndLN, or spleen (Extended Data Figure 6d). By UMAP analysis of the lungs, two new Treg clusters appeared during infection with differential expression of activation and differentiation markers (Extended Data Figures 8a and 8b). Using Ki-67 expression to separate proliferating cells, we observed an expansion in the frequency of Vβ9^+^ Tregs in the lungs that were enriched in the eTreg_1 cluster (Extended Data Figures 8c-8e). Together, these results indicate that our mass cytometric approach is capable of identifying CD8^+^ and conventional CD4^+^ T cell populations specific for endogenous viral antigens across different tissues. In addition, we identified an expansion of a Treg clonal population in response to flu infection.

### Expanded T cells using different Vβ chains adopt distinct differentiation states in response to immunization versus primary infection with influenza

We next asked whether prophylactic vaccination of mice would change the clonality or differentiation state of responding T cells in response to flu infection. We vaccinated mice by intramuscular (IM) injection of live flu virus (PR8). We then allowed mice to generate memory cells for one month prior to rechallenging mice with a respiratory flu infection (Figure 5a). Clustering analysis was performed to analyze different T cell populations in the context of no vaccination, IM vaccination, primary flu-infection, and flu rechallenge after prior IM vaccination. There was no difference in the composition of CD8^+^ T cell populations at baseline between vaccinated mice or naïve mice (Figure 5b). However, in the context of primary infection, there was a large representation of T_eff_ populations, similar to our previous analysis (Figure 4). However, in flu-challenged mice that were previously vaccinated, there was a dominant population present that expressed CD69 and CD39, which was not present in other memory populations (Figures 5b, 5c). Their expression of CD69, but not other markers of recently-activated effector cells such as T-bet, KLRG1, and Granzyme B, is consistent with a CD8^+^ tissue-resident memory (T_rm_) cell phenotype^47^. Employing PD-1 and Ki-67 as pan-activation markers for all CD8^+^ T cells, we observed a substantial expansion of only the Vβ8.3^+^ CD8^+^ T cells in mice that were rechallenged after vaccination (Figures 5d and 5e). In contrast, we did not see an expansion of Vβ6^+^ or Vβ7^+^ CD8^+^ T cells in rechallenged mice as was seen in mice with primary flu infection (Figures 5e and 4f-4g). Next, we assessed the activation and differentiation state of the Vβ8.3^+^ CD8^+^ T cells in the primary infection versus rechallenge by visualizing Vβ8.3^+^ cells by UMAP (Figure 5f). Interestingly, the Vβ8.3^+^ CD8^+^ T cells were enriched in the T_eff_ states in the primary infection, but in rechallenged mice, they were instead enriched in the T_rm_ state (Figure 5f). The exclusive enrichment of Vβ8.3^+^ CD8^+^ T cells in rechallenged mice within the T_rm_ cluster may indicate that vaccination can change the differentiation state of flu-specific cells that are recalled in response to respiratory flu infection. Furthermore, the Vβ6^+^ and Vβ7^+^ CD8^+^ T cells that were enriched in T_eff_ states in the primary infection did not expand with rechallenge, indicative of changes in the clonal dominance of T cells after vaccination. Our results also suggest that CD8^+^ T cells with different specificities may differentiate asymmetrically into different types of memory cells after vaccination, resulting in restricted secondary T cell specificities against a respiratory flu infection.

**Figure 5.**
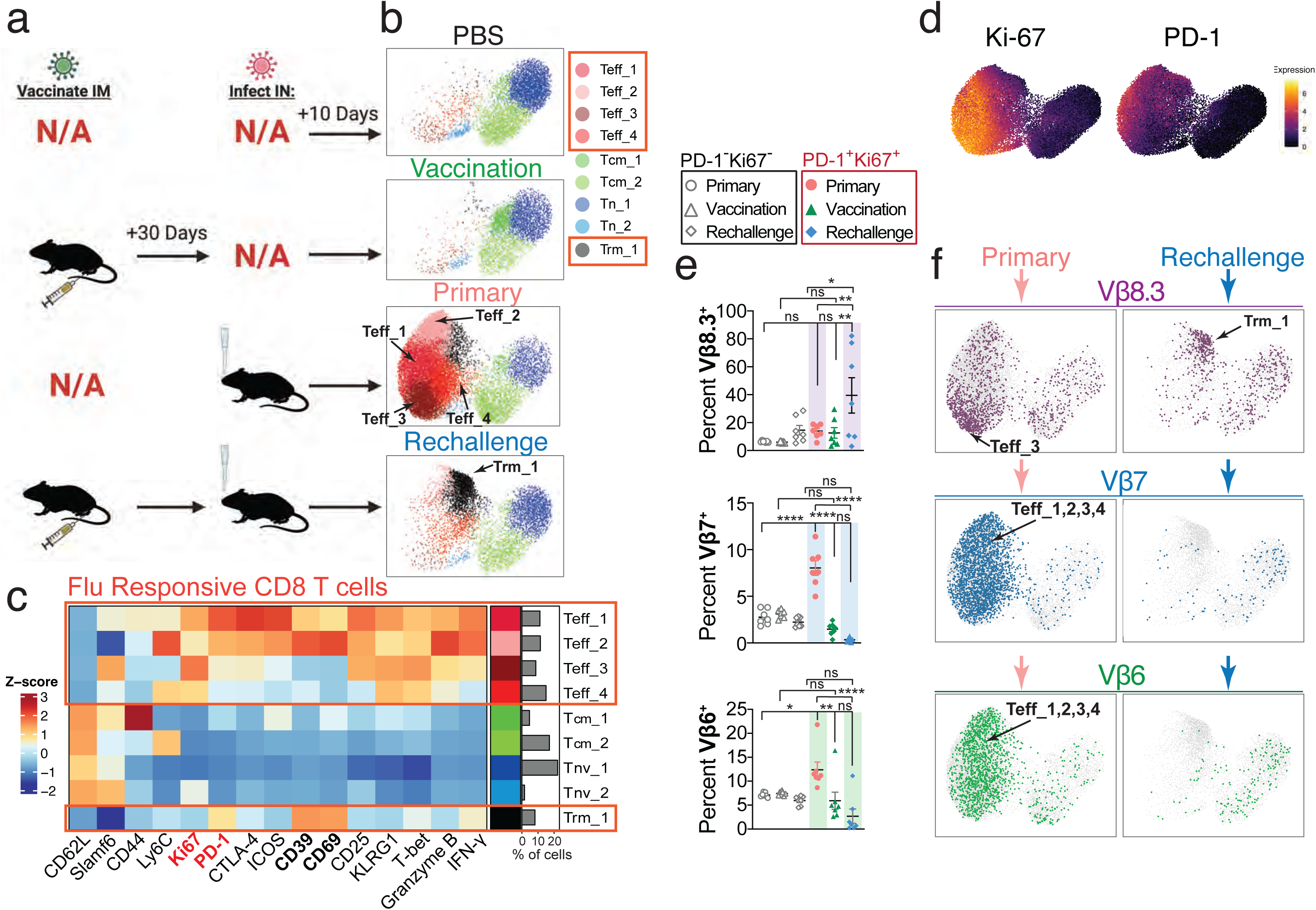
| Vβ8.3+ CD8s have different phenotypes and frequencies in the context of influenza immunization model. (a) C57BL/6J mice were vaccinated intramuscularly (IM) and later rechallenged intranasally (IN), vaccinated IM but not rechallenged, challenged with flu (IN) without prior vaccination, or injected with a PBS vehicle control. (b) UMAP visualization of CD8^+^ T cells based on expression of non-TCR proteins. (c) Heatmap of non-TCR protein expression annotated by cluster and fraction of cells falling into each cluster. (d) UMAP visualization of CD8^+^ T cells colored by the expression Ki-67 and PD-1 from all cohorts of mice. (e) Comparison of the frequencies of CD8^+^ T cells using Vβ8.3, Vβ7, and Vβ6 in non-proliferating (PD-1^−^Ki-67^−^) and proliferating (PD-1^+^Ki-67^+^) cells in the lungs of primary, vaccinated, or rechallenged mice. (f) UMAP visualization of pooled CD8^+^ T cells from primary or rechallenged mice colored by the expression of Vβ8.3, Vβ7, and Vβ6. Results from n=7 for PBS, n=7 for Primary, n=7 for Vaccination, n=7 for Rechallenge *P<0.05, **P<0.01, ***P<0.001 by one-way ANOVA, mean ± s.e.m.

In parallel, we also examined the impact of vaccination on the clonal behavior of conventional CD4^+^ T cells and Tregs. Like CD8^+^ T cells, conventional CD4^+^ T cells also contained a CD69^+^CD39^+^ population that expanded specifically in mice that were rechallenged with flu after vaccination (Extended Data Figures 9a and 9b). Analysis of the proliferating pool (Ki-67^+^) of CD4^+^ conventional T cells (Extended Data Figure 9c) revealed Vβ14^+^ cells that expanded in flu-infected mice independent of vaccination status (Extended Data Figures 9d and 7d). However, in the flu-infected mice without prior vaccination, Vβ14^+^ CD4^+^ conventional T cells were enriched in Teff_1 and Tem_2 clusters, while in vaccinated mice rechallenged with flu, they were instead enriched in the T_rm_ cluster, paralleling the differences observed in Vβ8.3^+^ CD8^+^ T cells between these contexts (Figure 6f). Examination of the Treg compartment showed that the Vβ9^+^ Tregs that expanded during primary flu infection, did not expand in pre-vaccinated mice (Extended Data Figure 10). Taken together, these results demonstrate that vaccination prior to respiratory flu infection alters the CD8^+^, CD4^+^, and Treg TCR repertoire and the differentiation state of cells expressing specific TCRs during a subsequent flu infection.

**Figure 6.**
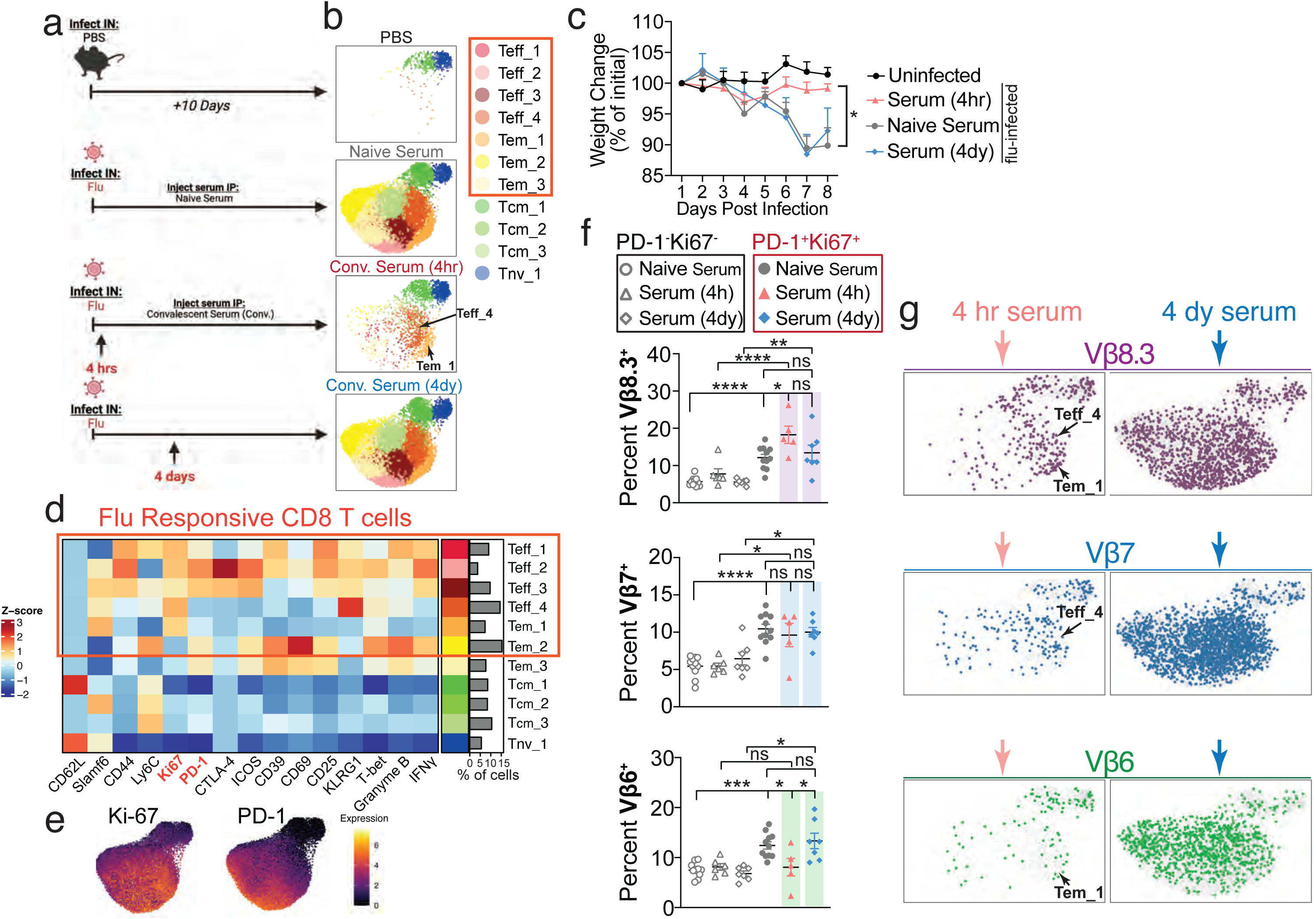
| Early convalescent therapy restricts the CD8^+^ T cell repertoire during influenza infection. (a) C57BL/6J mice were infected with influenza IN and then either untreated or treated with convalescent serum at 4 hours or 4 days after infection, and lungs were analyzed at day 10 for the T cell response. (b) UMAP of CD8^+^ T cells based on the expression of non-TCR proteins. (c) Weight change as percent of initial weight of uninfected and flu-infected mice treated with serum as indicated. (d) Heatmap of non-TCR protein expression annotated by cluster and fraction of cells falling into each cluster. (e) UMAP visualization of CD8^+^ T cells colored by the expression Ki-67 from all cohorts of mice. (f) Comparison of the frequencies of CD8^+^ T cells using Vβ8.3, Vβ7, and Vβ6 in non-proliferating (PD-1^−^Ki-67^−^) and proliferating (PD-1^+^Ki-67^+^) cells in the lungs of flu-infected mice treated with naive or convalescent serum at specified times after infection. (g) UMAP visualization of pooled CD8^+^ T cells from early vs. late convalescent serum treated mice colored by the expression of Vβ8.3, Vβ7, and Vβ6. Results from n=7 for PBS, n=14 for naïve serum-treated infected mice, n=7 for convalescent serum-treated infected mice at 4 hours, n=7 for convalescent serum-treated infected mice at 4 days, *P<0.05, **P<0.01, ***P<0.001 by one-way ANOVA for CyTOF analysis and two-way ANOVA for weight curves, mean ± s.e.m.

### Transfer of convalescent serum shortly after respiratory influenza infection mimics aspects of prophylactic vaccination by reshaping the TCR repertoire of responding T cells

Our prior results indicated that immunological memory alters the TCR repertoire and functional state of responding T cells after flu rechallenge. These changes could result from the formation of memory T cells with flu-specific TCRs. Alternatively, altered T cell responses could be due to T cell-extrinsic factors, such as antibody responses that could support or prevent T cell priming by antigen presenting cells or impact the viral load and thus the magnitude of the T cell response. Therefore, we investigated the effect of antibodies against flu on the T cell response by treating mice with convalescent serum from mice that recovered from respiratory flu infection. This approach also has therapeutic implications, as convalescent serum therapy is a standard form of treatment for severe respiratory viral infections, such as COVID-19, but its impact on the antiviral T cell response remains incompletely understood^48^.

We therefore tested whether the administration of serum therapy early after infection (4 hours post-infection) or just prior to the development of disease symptoms (4 days post-infection) impacted the T cell response to influenza infection (Figure 6a). We observed that mice were protected from weight loss when treated early during infection (Figure 6c). Mice receiving serum early in infection exhibited effector and memory CD8^+^ T cell populations that expressed lower levels of inflammatory markers, though they still expressed general activation markers such as PD-1 and Ki-67 (Figures 6d, 6e). When comparing distinct clonal populations within these proliferating CD8^+^ T cells, mice receiving serum early during infection had augmented frequencies of Vβ8.3^+^ CD8^+^ T cells, while an expansion in Vβ6^+^ cells was no longer present (Figure 6f), similar to our observations in response to vaccination (Figure 5). However, these Vβ8.3^+^ clonal populations are preferentially clustered in a Tem_1 population, as in the primary infection (Figure 6g), and not toward a T_rm_ phenotype, as observed after vaccination and rechallenge (Figures 6g and 5f). We hypothesize that this change in differentiation states occurs because memory T cell differentiation requires time to develop after antigen encounter. In contrast, the Vβ7^+^ CD8^+^ T cell response was unaltered by serum transfer, and CD8^+^ T cells expanded and differentiated equivalently to what was observed with primary flu infection (Figures 6f and 6g). These results indicate that anti-flu antibodies can impact the CD8^+^ T cell response to infection, both in the specificity and the differentiation state of responding CD8^+^ T cells.

T cell priming occurs early during an infection^49,50^. Therefore, we hypothesized that treatment with convalescent serum later in an infection, but prior to the onset of severe symptoms (i.e., weight loss), might have a minimal impact on the T cell repertoire and differentiation states. Consistent with this hypothesis, we did not observe any protection from weight loss in mice treated with convalescent serum 4 days after infection (Figure 6c). We also did not observe any changes in the differentiation of CD8^+^ T cells using Vβ8.3, Vβ6, or Vβ7 TCR chains, all of which similarly expanded and differentiated into multiple T_eff_ and T_em_ phenotypes, compared to mice undergoing a primary flu infection (Figure 6b). In contrast to early serum transfer, serum treatment later in the infection resulted in equivalent expansion and differentiation of Vβ8.3^+^ CD8^+^ T cells as seen in the primary infection (Figures 6f and 6g). Thus, antibody responses to flu, particularly in the context of transferred antibodies from vaccinated mice that have affinity matured, class switched, and likely have neutralizing capacity, altered the TCR specificity of responding CD8^+^ T cells as well as their differentiation program only when administered early after flu infection.

In contrast to the impact on CD8^+^ T cells, mice receiving serum exhibited fewer changes in CD4^+^ T cells (Extended Data Figures 11a-11c). While there were noticeable reductions in the generation of T_eff_ populations with serum treatment at 4 hours post-infection, they were mild reductions. Furthermore, there were not any changes in the frequency of CD4^+^ T cells using the Vβ14 chains with serum treatment early or late after infection, although early serum transfer had modest impacts on the differentiation state of these Vβ14^+^ CD4^+^ T cells (Extended Data Figures 11d and 11e). Interestingly, however, we did find that the expansion of Vβ9^+^ Tregs was blunted by convalescent serum transfer 4 hours post-infection (Extended Data Figure 12), which matched the impact of IM vaccination (Extended Data Figure 10). This result, in combination with our observations in the vaccination model, suggests that neutralizing antibodies are sufficient to prevent the expansion of this Vβ9^+^ Treg population even in the absence of memory T cell responses.

## Discussion

The T cell receptor (TCR) repertoire changes dramatically as a result of the clonal expansion of antigen-specific T cells during immune responses to pathogens and cancer. However, it is not fully understood to what extent this repertoire must be remodeled to achieve a successful immune response or therapeutic intervention. This highlights the need for better systems to track the clonality and phenotype of T cells during immune responses. Here we demonstrated that mass cytometry (CyTOF) can be used to track T cell populations with specific Vβ and Vα chains, allowing us to identify several antigen-responsive T cell populations while also assessing their functionality, differentiation state, and proliferation across different disease models.

This approach has complementary strengths and weaknesses in comparison to single-cell RNA-sequencing (scRNA-seq) that is paired with TCR sequencing (Extended Data Figure 5). For paired scRNA-TCR sequencing, where ∼45% of TCR Vβ chains were not assigned, TCR assignment can be affected by transcript recovery, leading to a variable proportion of input cells lacking full TCR alpha-beta assignment. In contrast, while our CyTOF method only missed assigning ∼25% of TCR Vβ chains, the approach is currently limited by the lack of available antibodies against other TCR variable chains (i.e., Vβ1, 15, 16, 18, 20). In addition, the antibodies against Vβ5.1/2 (BD clone MR9-4) and Vβ8.1/2 (BD clone MR5-2) each bind to two Vβ chains that cannot be distinguished by CyTOF but can be deconvoluted by sequencing. For these reasons, both datasets were not fully overlapping, but this was anticipated as experiments were performed in distinct mouse cohorts at two different laboratories. Therefore, it was impressive that despite these limitations and differences in methods, there were comparable frequencies of Vβ chain assignments broadly and specifically when assessing SIINFEKL-reactive CD8^+^ T cells (Extended Data Figure 5i). CyTOF could be used as a high-throughput pre-screening tool to narrow the cell sorting and higher cost scRNA-seq platforms to specific Vα and Vβ and chain-expressing T cells^511^.

We validated our CyTOF method using OVA as a model antigen to first identify an endogenous OVA-specific CD8^+^ T cell population and examine its differentiation and function in different disease contexts. We found that OVA-specific CD8^+^ T cell populations were enriched within Vβ14^+^ CD8^+^ T cells in the *LADD-OVA* infection model and two OVA-expressing tumor models. Further, we identified *Listeria* antigen-specific CD8^+^ T cells and conventional CD4^+^ cells and validated the specificity of these populations using peptide restimulation, tetramer-based assays, and scRNA-seq data. Interestingly, our system was sensitive enough to identify changes in both CD4^+^ conventional T cells and CD8^+^ T cell subtypes that recognized endogenous *Listeria*-derived antigens, even when the strong model antigen OVA was present. We detected infiltration of OVA-specific Vβ14^+^ CD8^+^ T cells in B16F10 tumors that expressed OVA and expansion of these Vβ14^+^ CD8^+^ T cells in the tumor-draining lymph nodes. Thus, our CyTOF approach can capture the behavior of T cell populations across different disease models by measuring changes in the expansion of specific Vβ chain expressing T cells, even in the context of an immunodominant antigen such as OVA.

A major finding using our CyTOF approach during respiratory influenza infection was the enrichment of CD8^+^ T cells with a T_rm_-like phenotype and the restriction of the anti-influenza TCR repertoire (Vβ8.3^+^ enriched) with vaccination or convalescent serum therapy. Our data also suggests that influenza vaccination improves the immune response to subsequent infection by driving a subset of Vβ8.3^+^ CD8^+^ T cells to take on a tissue resident memory-like differentiation state. Tissue resident memory cells allow for control of broad viral serotypes against several viruses ranging from influenza to SARS-COV-2^52^. The Vβ8.3^+^ T_rm_-like cells were the most expanded population during infection in mice that were previously vaccinated. This suggests that this population of CD8^+^ T cells expressing the Vβ8.3 chain are poised to become memory cells, while populations expressing other Vβ chains are either less efficient at differentiating toward a T_rm_ phenotype or are outcompeted by Vβ8.3^+^ T_rm_-like cells upon rechallenge. Such differences in differentiation capacities have been observed previously for effector CD8^+^ T cells during respiratory infections, where memory cell selection was driven by TCR avidity for peptide-MHC^53^. Further studies must be conducted to determine if this Vβ8.3^+^ clonal population is critical for improved protection from disease upon rechallenge with the same serotype, and potentially against different serotypes. More broadly, it should be investigated why this particular clonal population is favored to differentiate toward a tissue resident memory-like state.

Convalescent serum therapy has been utilized to treat several respiratory viral infections due to the presence of neutralizing antibodies that can limit viral spread and improve viral phagocytosis by APCs^54^. We were interested in analyzing the effect of this therapy on the clonal T cell response since it has been shown previously that convalescent plasma therapy from mild or severe COVID-19 patients can have differing effects on the percentage of activated effector T cells and memory T cells^55^. Furthermore, the timing of convalescent serum therapy has also been shown to correlate with its efficacy in preventing severe disease^56–61^. Indeed, treatment of mice with convalescent serum during the earliest stages of infection broadly restricted the expansion of inflammatory effector T cell populations in the lungs and reduced disease severity (Figure 6c). However, convalescent serum 4 hours after infection did lead to the increased expansion of the Vβ8.3^+^ CD8^+^ T cells, which exhibited an effector memory phenotype. This may indicate that very early convalescent serum therapies directly impact viral load through antibody activities, but also impact the activation and differentiation of specific CD8^+^ T cell clones against the virus. The impact of antiviral antibody on T cell differentiation has been hypothesized previously and attributed to various effects of the presence of antibodies early, such as reducing the levels of circulating inflammatory cytokines, which could impact T cell response and disease severity, or antibody-mediated presentation of antigen that leads to expansion of T cells with particular specificities^59,60^. However, it is challenging to draw direct parallels between observations in patients or other animal models with different respiratory viruses due to the difference in viruses, the physiology of infection and immune responses in each species, and the delivery and dosing of virus in each experimental model^62,63^. Nevertheless, our results do concur with a series of studies that clearly show that earlier antibody treatment is more effective in controlling severe respiratory viral disease. We show that early treatment does impact the repertoire and differentiation of specific T cells, and thus our model will provide a new system to dissect whether altered T cell responses with convalescent serum therapy contribute to reduced disease severity.

Importantly, an advantage of this CyTOF approach is its capacity to immediately be translated to the setting of human diseases. Human Vα and Vβ chain specific antibodies are commercially available, making it possible to identify clonal T cell responses to chronic infections, new pathogens, microbiota, and cancer in humans^64–66^. Additionally, we found that this approach can identify Vβ chain expressing Tregs that expand during infection, which has been difficult to do historically^67^. This approach can also be used to rapidly identify antigens in several human diseases to inform the development of vaccines to treat cancers or infections. This would complement the use of strategies such as CITE-seq and scTCR-seq, which have been used recently to understand the behavior of clones in cancer patients in response to checkpoint blockade immunotherapy^68,69^. Importantly, our CyTOF approach can be used as a preliminary screening method for identifying T cell populations of interest, independent of the identity of their antigenic target, prior to TCR sequencing of populations enriched for expansion during an immune response^31^. The scalability of our CyTOF approach should allow for broader analyses across tissues, time points, or therapeutic perturbations prior to TCR sequencing experiments, vastly improving sequencing efficiency and cost.

In summary, we report a new application of CyTOF to quickly identify and phenotype clonal T cell populations across several disease models. In addition, we used these common Vβ and Vα chains to track the states of particular clonal populations of interest during different therapeutic contexts, such as vaccinations and convalescent serum therapy for respiratory infection with flu. Collectively, our study provides a new approach for identifying and phenotyping relevant T cell populations containing clones specific for antigens across multiple disease models that can be used with the aim of improving the efficacy of immunotherapies.

## Methods

### Animals

C57BL/6J wildtype mice were obtained from Jackson laboratories (JAX:000664). For tumor studies, syngeneic C57BL/6J mice were inoculated with 5.0×10^5^ MC38, 5.0×10^5^ MC38 cells engineered to express ovalbumin (OVA), 5.0×10^5^ B16F10 cells, or 5.0×10^5^ B16F10 cells engineered to express OVA in PBS subcutaneously. All the experiments were conducted according to the Institutional Animal Care and Use Committee guidelines of the University of California, Berkeley.

### Cell lines

MC38, MC38-OVA, B16F10, and B16F10-OVA were kindly provided by Dr. Jeff Bluestone’s lab^40^. All cell lines were maintained in DMEM (GIBCO) supplemented with 10% FBS, sodium pyruvate (GIBCO), 10mM HEPES (GIBCO), and penicillin-streptomycin (GIBCO). Tumor cells were grown at 37℃ with 5% CO_2_.

### Intravenous *Listeria* (LADD) infection

All strains of *L. monocytogenes* used in this study were a gift from the Portnoy laboratory at University of California, Berkeley and were derived from the wild type 10403S strain. The LADD constructs were based on Lm ΔactA/ΔinlB. LADD-OVA expresses a secreted ActA-OVA fusion as described under the control of the actA promoter^70^. All strains were cultured in filter-sterilized nutrient-rich Brain Heart Infusion (BHI) media (BD Biosciences) containing 200 μg/mL streptomycin (Sigma-Aldrich). Overnight cultures were grown in BHI + 200 μg/mL streptomycin at 30℃. The following day, bacteria were grown to logarithmic phase by diluting the overnight in new BHI + 200 μg/mL streptomycin and culturing at 37℃ shaking. Log-phase bacteria were washed and frozen down in 9% glycerol/PBS. For infections, frozen stocks were diluted in PBS to infect via the tail vein with 1 x 10^7^ CFU log-phase bacteria. The mice were euthanized 5 days post infection, and the spleen was collected for flow analysis.

### Intranasal Influenza Infection and intramuscular vaccination

A stock of PR8 H1N1 influenza was a gift from the Arpaia laboratory at Columbia University. Prior to treatment of mice, aliquots were thawed and diluted in sterile PBS such that the desired dose was 54 PFU. Mice were then anesthetized with isoflurane and treated intranasally. For vaccination studies, viral stocks were diluted to inject 540 PFU intramuscularly into the right thigh. Mice were left for a month for viral clearance and generation of memory response before continuing studies.

### Convalescent serum treatment

Mice were infected with 54 PFU intranasally and left for a month for viral clearance and generation IgG. Serum was harvested by collecting blood via cardiac puncture and incubating blood at room-temperature to allow coagulation. Samples were centrifuged aliquoted and maintained at 4℃. Quantification of IgG for samples was done by diluting serum samples and using the Mouse IgG ELISA commercial kit (Molecular Innovations). For treatment, infected and control mice were given a single dose of 50μg/mL serum derived from naive or previously infected mice by intraperitoneal (i.p.) injection.

### Tissue Collection and preparation for Flow cytometry

Flow cytometry was performed on an BD LSR Fortessa X20 (BD Biosciences) or LSRFortessa (BD Biosciences) and datasets were analyzed using FlowJo software (Tree Star). Single cell suspensions were prepared in ice-cold FACS buffer (PBS with 2mM EDTA and 1% BS) and subjected to red blood cell lysis using ACK buffer (150mM NH4Cl, 10mM KHCO3, 0.1mM Na2EDTA, pH7.3). Dead cells were stained with Live/Dead Fixable Blue or Aqua Dead Cell Stain kit (Molecular Probes) in PBS at 4℃. Cell surface antigens were stained at 4℃ using a mixture of fluorophore-conjugated antibodies. Surface marker stains for murine samples were carried out with anti-mouse CD3 (17A2, BioLegend), anti-mouse CD4 (RM4-5, BioLegend), anti-mouse CD8a (53-6.7, BioLegend), anti-mouse, CD44 (IM7, BioLeged), anti-mouse CD45 (30-F11, BioLegend), anti-H-2Kb MuLV p15E Tetramer-KSPWFTTL (MBL), anti-H-Kb-A2/SIINFEKEL tetramer (NIH tetramer core), anti-IAb/NEKYAQAYPNVS tetramer (NIH tetramer core) in PBS, 0.5% BSA. Cells were fixed using the eBioscience Foxp3/Transcription Factor staining buffer set eBioscience), prior to intracellular staining. Intracellular staining was performed using anti-mouse Foxp3 (FJK-16S, eBioscience), anti-mouse TNF-α (MP6-XT22, BioLegend), anti-mouse IFN-γ (XMG1.2, eBioscience), at 4℃, according to manufacturer’s instructions. Cells were resuspended in PBS and filtered through a 70-μm nylon mesh before data acquisition. Datasets were analyzed using FlowJo software (Tree Star).

### Peptide Restimulation Assays

Resected tumors and lung tissues were minced to 1 mm^3^ fragments and digested in RPMI media supplemented with 4-(2-hydroxyethyl)-1-piperazi-neethanesulfonic acid (HEPES), 20 mg/mL DNase I (Roche), and 125 U/mL collagenase D (Roche) using an orbital shaker at 37℃. Cells from lymphoid organs were prepared by mechanical disruption pressing against a 70-μm nylon mesh. All the cell suspensions were passed through 40 μm filters before in vitro stimulation. Cytokine staining was performed with 3-5×10^6^ cells in Opti-MEM media supplemented with Brefeldin A (eBioscience), 1μg/mL peptides, or 10 ng/mL phorbol 12-myristate 13-acetate (PMA) (Sigma), and 0.25 μM ionomycin (Sigma). Fixation/permeabilization of cells was conducted for intracellular staining using the eBioscience Foxp3 fixation/permeabilization kit (BioLegend) or Tonbo Foxp3 / Transcription Factor Staining Buffer Kit.

### Tissue collection and preparation for Mass Cytometry

To collect tissues for analysis, at the experimental endpoint mice were euthanized with CO_2_ inhalation. Mice were then tracheally perfused with PBS + 5mM EDTA, and relevant organs were collected. When tumor or lung tissue was collected, these tissues were finely minced and then digested at 37℃ in digestion buffer: RPMI 1640 (UCSF Media Production) with 1 mg/mL collagenase IV (Worthington Biochemical) and 0.1 mg/mL DNase I (Sigma-Aldrich). All organs were then processed into a single cell suspension over a 70uM filter and washed with PBS + 5mM EDTA. Cells were resuspended 1:1 with PBS + 5mM EDTA and 100μM Cisplatin (Sigma-Aldrich) (diluted in PBS + 5mM EDTA) for 60s before quenching with 1:1 with PBS + 0.5% BSA + 5mM EDTA to determine viability^27^. Cells were again centrifuged and resuspended in PBS + 0.5% BSA + 5mM EDTA. Samples were aliquoted into individual cluster tubes at ∼1-10×10^6^ cells per tube and then fixed at 1.6% PFA (Thermo Scientific). After quenching the fixation with PBS + 0.5% BSA + 5mM EDTA and washing out the fixative, cells were frozen at –80℃ for subsequent CyTOF analysis.

### Mass Cytometry Antibodies

All antibodies used in this study can be found in Extended Data Table 1. Antibodies were purchased unlabeled and conjugated to heavy metals in-house^71^. Antibody conjugation to heavy metal tags was done using the MaxPar Antibody Conjugation Kit (Fluidigm) according to the manufacturer’s protocol. After labeling, antibodies were diluted to 0.2mg-0.5mg/mL in antibody stabilization solution (Candor Bioscience) and stored at 4℃ until use. Before using experimentally, conjugated antibodies were titrated on mouse tissue to determine optimal staining concentration (Extended Data Table 1). Prior to sample staining, all antibodies were pooled together into either surface or intracellular master mixes and stained together.

### Cellular Barcoding

Mass tag cellular barcoding was performed as previously described^72^. Briefly, 1×10^6^ cells from a set of 20 samples were stained with a unique ‘barcode’ that combines 3 Pd isotypes out of 6 total isotypes. This stain was done in 0.02% saponin (Fluidigm’s 10× Barcode Perm Buffer diluted in PBS). These 20 samples were then pooled together into 1 tube, washed with PBS + 0.5% BSA + 0.02% NaN_3_, and then all ∼20×10^6^ cells were stained together for CyTOF. After running these samples by CyTOF, sample data were then deconvoluted according to their Pd isotype barcode as previously described^72^.

### Staining for CyTOF

After cellular barcoding, each sample was aspirated to exactly 95μL of volume. 5uL of TruStain FcX (anti-mouse CD16/32) (Biolegend) was then added and incubated at room temperature for 5 minutes. After Fc block, 400μL of the surface staining master mix was added to the samples for a total staining volume of 500μL. Sample tubes were then incubated for 30 minutes at room temperature shaking and washed with PBS + 0.5% BSA + 0.02% NaN_3_. Each sample was then permeabilized with 100% Methanol and incubated at 4°C. Samples were then quenched and washed twice with PBS + 0.5% BSA + 0.02% NaN_3_. Intracellular master mix was then added to each sample for a final staining volume of 500μL and incubated for 30 minutes at room temperature shaking. Following the intracellular stain, samples were washed with PBS + 0.5% BSA + 0.02% NaN_3_ and resuspended in 1mL PBS + 2-4% PFA + 1:4000 191/193Ir DNA intercalator (Fluidigm), and left to stain overnight or up to 7 days at 4°C^73^.

### CyTOF data collection and normalization

Immediately prior to data acquisition on the instrument, the sample of interest was removed from 4°C, washed with ddH20, and then washed with Cell Acquisition Solution (CAS) (Fluidigm). For sample running, a bead normalization solution was made by diluting Calibration beads, EQ™ Four Element (Fluidigm), 1:50 by volume in CAS. The sample was then resuspended in 1mL of this bead solution, counted, and diluted to 1×10^6^ cells/mL prior to running on a CyTOF 2 mass cytometer. The entirety of each sample was then collected. After data collection and FCS file creation, samples were normalized as previously described using the internal bead control to account for acquisition fluctuations over time^73^.

### Semi-supervised CyTOF clustering and TCR chain analysis

After cellular debarcoding and normalization of samples, individual fcs files were uploaded to Cell Engine. For downstream analysis, T cells were gated according to the following strategy: live (cisplatin^−^, 191/193Ir^+^), CD45^+^, CD3^+^. From there, CD8 T cells were then gated as CD8^+^, conventional CD4^+^ T cells were gated as CD4^+^ FoxP3^−^, and Tregs were gated as CD4^+^ FoxP3^+^. These 3 populations were then exported from Cell Engine as fcs files for each sample for each tissue for each experiment. These fcs files were then imported into R, arcsinh transformed, and compiled into a master data frame for use in the semi-supervised clonotyping assignment script as follows:

First, a matrix is created to store threshold assignments for each Vα and Vβ chain. The default threshold set is an arcsinh-transformed value of 1.5 (arcsinh(x/5) >= 1.5), but this can be changed by the used based on the staining pattern of a particular antibody (Extended Data Figure 13a, 13b). Then, new columns are created in the master dataframe for each subsequent Vα and Vβ assignments to be written in. Upon running the master function, each cell in the data frame will be iterated through and assessed for the expression value of each individual Vα and Vβ chain. If there is no Vα channel that has a value over the threshold value, the cell’s Vα column will be filled in with ‘None.’ The same process is implemented for the Vβ channels. If there is more than one Vα that has a value over the threshold value, the cell’s Vα column will be filled in with ‘Unassigned.’ The same process is implemented for the Vβ channels. If the cell has exactly one Vα channel that has a value over the threshold value, the cell’s Vα column will be populated with that channel name. The same process is implemented for the Vβ channels. After iterating through each cell in the dataframe, the function creates an additional column called ‘Clonotype’ and concatenates the Vα and Vβ columns into this column (ie, Vα2_Vβ14). Finally, the master data frame is written as a .csv file into the working directory such that clonotype assignment information is maintained for use in downstream analyses.

Rphenograph was used for clustering, and separate experiments, organs, and cell populations (i.e., CD8s, CD4s, Tregs) were clustered separately. Clustering was done in an unbiased manner using the following parameters: neighbors = 100, mindist = 0.4, k_pheno = 100. An analysis of this computational approach is included in Extended Data Figure 13. Extended Data Fig. Panels 13c (CD8^+^ T cells) and 13f (CD4^+^ T cells) show the percentage of cells categorized as ‘Assigned,’ ‘Unassigned,’ or ‘None, by the semi-supervised script for each experiment. For the ‘Unassigned’ condition, cells are further categorized by whether the Vα, Vβ, or both chains were ‘Unassigned.’ This is further demonstrated at the individual chain level, with Extended Data Figures panels 13d,13g showing the Vβ assignment for CD8^+^ and CD4^+^ T cells, and Extended Data Figures 13e, 13h showing the Vα assignment for CD8^+^ and CD4^+^ T cells.

## Acknowledgements

We acknowledge the UCSF Parnassus Flow Cytometry CoLab (RRID:*SCR_018206*) for assisting in generating mass cytometry data and the UC Berkeley Cancer Research Laboratory Flow Cytometry Facility. We also thank D. Portnoy (University of California, Berkley) for providing attenuated *Listeria* strains for this study. Research reported here was supported in part by Grants from the NIH 1DP2CA247830-01, P30 DK063720, and the NIH S10 Instrumentation Grant S10 1S10OD018040-01. J.G.C. is a HHMI Gilliam Fellow. M.D. is a Pew-Stewart Scholar and a St. Baldrick’s Scholar with generous support from Hope with Hazel.

## Data availability

Additional data available upon request.

## Code availability

Automated Clonotyping assignments script is available at Github: *<u>github.com/SpitzerLab/Semi-</u> <u>supervised-Clonotyping-Assignments</u>*

## Extended Data Figure Legends

**Extended Data Figure 1 | Tracking a *Listeria*-specific CD8^+^ T cell response against *LADD* by TCR use and phenotyping with CyTOF.** (a) Gating strategy for all CyTOF analyses preceding input into semi-supervised clonotype assignment script. (b) Representative flow plots and quantification of proliferating CD8^+^ T cells from spleens 5 days after *LADD, LADD-OVA, or PBS* injection IV. (c) Bar plot of Vα_2, Vβ_14, and Vα_2_Vβ_14 percentages in each cluster for conditions *LAAD* and *LAAD-OVA*. (d) UMAP visualization of pooled CD8^+^ T cells colored by the expression of Vβ12^+^ in *LADD*, *LADD-OVA*, or PBS infected mice. For all plots, **P<0.05, **P<0.01, ***P<0.001 by unpaired two-tailed student’s T-test, mean ± s.e.m.

**Extended Data Figure 2 | Tracking *Listeria*-specific CD4^+^ T cells against *LADD* by TCR use and phenotyping with CyTOF.** (a) Representative flow plots and quantification of proliferating (Ki-67^+^) CD4^+^ T cells from spleen 5 days after *LADD, LADD-OVA, or PBS* injection. (b) UMAP visualization of CD4^+^ T conventional (non-Treg) cell clusters based on expression of non-TCR proteins. (c) Heatmap of non-TCR protein expression annotated by cluster and fraction of cells falling into each cluster. (d) UMAP visualization of CD4^+^ T cells colored by the expression Ki-67 in *LADD*-infected mice. (e) Frequency of Ki-67^+^ and Ki-67^−^ CD4^+^ T cells using specific TCR Vβ (left) or Vα chains (right) in response to PBS, *LADD*, or *LADD-OVA*. (f-h) Frequency of Vβ13^+^ (f), Vβ14^+^ (g), or Vβ13^+^Vα2^+^ among Ki-67^+^(h) versus Ki-67^−^ CD4^+^ T cells in *LADD* or *LADD-OVA* infected mice. (i-k) UMAP visualization of pooled CD4^+^ T cells colored by the expression of Vβ13^+^ (i), Vβ13^+^Vα2^+^ (j), and Vβ14 (k) in *LADD*, *LADD-OVA*, or PBS injected mice. (m) Bar plot of Vβ13^+^, Vβ13^+^Vα2^+^, and Vβ14^+^ percentages in Teff clusters. Results from n=5 for PBS, n=4 for *LADD*, n=4 for *LADD-OVA*, *P<0.05, **P<0.01, ***P<0.001 by unpaired two-tailed Student’s t-test, mean ± s.e.m.

**Extended Data Figure 3 | CD8 response to *LADD*– and OVA-derived epitopes.** (a) Manual gating strategy for all CD8 and CD4 Teff subsets. (b) FMO controls for all fluorescent-conjugated Vβ and Vα antibodies in naive splenocytes +/− stimulation with PMA/Ionomycin. (c) Frequency of IFN-y^+^ CD8s in splenocytes in unstimulated controls versus peptide conditions. (d) Representative flow plots of IFN-y^+^ CD8s versus respective Vβ chains in splenocytes stimulated with SIINFEKL peptide. (e) Percent of IFN-y^+^ for respective Vβ^+^ CD8s in splenocytes stimulated with SIINFEKL peptide. (f) Representative flow plots of IFN-y^+^ CD8s versus respective Vβ chains in splenocytes stimulated with LLO1 peptide. (g) Percent of IFN-y^+^ for respective Vβ^+^ CD8 in splenocytes stimulated with LLO1 peptide. Results from n=5 for PBS, n=4 for *LADD*, n=4 for *LADD-OVA*, *P<0.05, **P<0.01, ***P<0.001 by unpaired two-tailed Student’s t-test, mean ± s.e.m.

**Extended Data Figure 4 | CD4 response to *LADD-*derived epitopes.** (a) FMO controls for all fluorescent-conjugated Vβ and Vα antibodies in naive splenocytes +/− stimulation with PMA/Ionomycin. (b) Frequency of IFN-y^+^ CD4s in splenocytes in unstimulated controls versus peptide conditions. (c) Representative flow plots of IFN-y^+^ CD4s versus respective Vβ chains in splenocytes stimulated with LLO2 peptide. (d) Percent of IFN-y^+^ for respective Vβ^+^ CD4s in splenocytes stimulated with LLO2 peptide. (e) Representative flow plots of IFN-y^+^ CD4s versus respective Vβ chains in splenocytes stimulated with LLO3 peptide. (f) Percent of IFN-y^+^ for respective Vβ^+^ CD4s in splenocytes stimulated with LLO3 peptide. Results from n=5 for PBS, n=4 for *LADD*, n=4 for *LADD-OVA*, *P<0.05, **P<0.01, ***P<0.001 by unpaired two-tailed Student’s t-test, mean ± s.e.m.

**Extended Data Figure 5 | Comparative analysis of Vβ14+Vα2+ CD8 expansion with scRNAseq during Lm infection.** *(a)* Schematic comparison of approach used in this study compared to Straub et al. (2023). (b) Re-annotated clusters identified by Straub et al. plotted in UMAP space. (c) Scaled gene expression within the scRNA-seq dataset of genes encoding the proteins measured in Fig. 1, stratified by cluster. (d) UMAP visualization of all clustered cells colored by Ki67 or PD-1 gene expression. (e) UMAP visualization of all clustered cells colored by TRBV31 assignment or TRAV14D-2 assignment (**left**), and UMAP visualization of TRBV31 assigned cells colored by TRAV14D-2 assignment (**right**). (f) UMAP visualization of all clustered cells colored by identified specificity. (g) Quantification of TRBV gene usage among SIINFEKL-reactive cells (right) or of cells of unidentified reactivity (left). (h) Quantification of TRBV gene usage among SIINFEKL-reactive cells expressing TRAV14D-2. (i) Quantification of TRBV/Vβ chain assignment of all CD8 T cells as measured by CyTOF or scRNA-seq. (j) Quantification of the TRBV/Vβ chains measured in the CyTOF assay of all CD8 T cells that were assigned a TRBV/Vβ chain by either CyTOF or scRNA-seq. (k) Spearman correlation analysis of data shown in (j).

**Extended Data Figure 6 | Expansion and activation of T cell subsets during flu infection across multiple tissues.** (a) Gating strategy for all CyTOF analyses preceding input into semi-supervised clonotype assignment script. (b) Representative flow plots and quantification of proliferating (PD-1^+^Ki67^+^) CD8^+^ T cells. (c) Representative flow plots and quantification of proliferating (Ki67^+^) Foxp3^−^ CD4^+^ Teff cells. (d) Representative flow plots and quantification of proliferating (Ki67^+^) Foxp3^+^ Tregs. For all plots, **P<0.05, **P<0.01, ***P<0.001 by one-way ANOVA, mean ± s.e.m.

**Extended Data Figure 7 | Identification of flu NP-specific Vβ14^+^ CD4^+^ T cells.** (a) UMAP visualization of CD4^+^ T conventional (non-Treg) cell clusters based on expression of non-TCR proteins. (b) Heatmap of non-TCR protein expression annotated by cluster and fraction of cells falling into each cluster. (c) UMAP visualization of CD4^+^ T cells colored by the expression Ki-67 in flu-infected mice. (d) Frequencies of Vβ14^+^ CD4 T cells in non-proliferating (Ki-67^−^) vs. proliferating (Ki-67^+^) cells across indicated tissues. (e) UMAP visualization of pooled CD4^+^ T cells colored by Vβ14 expression. (f) Quantification of fold-enrichment in Vβ-chain usage by CD4^+^ T cells stimulated with the PR8 NP peptide (LILRGSVAHKSCLPACV). (g) Schematic summarizing that Vβ14^+^ CD4^+^ T cells are enriched to recognize PR8 NP antigen from flu. Results from n=7 for PBS, n=7 for PR8, *P<0.05, **P<0.01, ***P<0.001 by unpaired two-tailed Student’s t-test, mean ± s.e.m.

**Extended Data Figure 8 | Identification and phenotyping of Vβ9+ Treg expansion after flu infection.** (a) UMAP of Treg populations based on expression of non-TCR proteins. (b) Heatmap of non-TCR protein expression annotated by cluster and fraction of cells falling into each cluster. (c) UMAP visualization of Tregs colored by the expression Ki-67 in flu-infected mice. (d) Frequencies of Vβ9^+^ Tregs in non-proliferating (Ki-67^−^) vs. proliferating (Ki-67^+^) cells across indicated tissues. (e) UMAP visualization of pooled Tregs from PBS or flu-infected mice colored by Vβ9 expression. Results from n=7 for PBS, n=7 for PR8, *P<0.05, **P<0.01, ***P<0.001 by unpaired two-tailed Student’s t-test, mean ± s.e.m.

**Extended Data Figure 9 | *Flu*-specific conventional CD4^+^ T cell differentiation, but not repertoire, is altered by vaccination.** (a) UMAP visualization of CD4^+^ T conventional (non-Treg) cell clusters based on expression of non-TCR proteins. (b) Heatmap of non-TCR protein expression annotated by cluster and fraction of cells falling into each cluster. (c) UMAP visualization of CD4^+^ T cells colored by the expression Ki-67 from all cohorts of mice. (d) Frequencies of Vβ14^+^ CD4 T cells in non-proliferating (Ki-67^−^) vs. proliferating (Ki-67^+^) cells in lungs of primary, vaccinated, or rechallenged mice. (e) UMAP visualization of pooled CD4^+^ T cells from primary or rechallenged mice colored by Vβ14 expression. Results from n=7 for PBS, n=7 for PR8, *P<0.05, **P<0.01, ***P<0.001 by one-way ANOVA, mean ± s.e.m.

**Extended Figure 10 | Vβ9+ Treg clonal population is only present in the setting of a primary infection.** (a) UMAP of Treg populations based on the expression of non-TCR proteins. (b) Heatmap of non-TCR protein expression annotated by cluster and fraction of cells falling into each cluster. (c) UMAP visualization of Tregs colored by the expression Ki-67 from all cohorts of mice. (d) Frequencies of Vβ9^+^ Tregs in non-proliferating (Ki-67^−^) vs. proliferating (Ki-67^+^) cells in lungs of primary, vaccinated, or rechallenged mice. (e) UMAP visualization of pooled Tregs from primary or rechallenged mice colored by Vβ9 expression. Results from n=7 for PBS, n=7 for PR8, *P<0.05, **P<0.01, ***P<0.001 by one-way ANOVA, mean ± s.e.m.

**Extended Data Figure 11 | Convalescent therapy has a limited effect on the CD4^+^ Teff repertoire during influenza infection.** (a) UMAP visualization of CD4^+^ T conventional (non-Treg) cell clusters based on expression of non-TCR proteins. (b) Heatmap of non-TCR protein expression annotated by cluster and fraction of cells falling into each cluster. (c) UMAP visualization of CD4^+^ T cells colored by the expression Ki-67 from all cohorts of mice. (d) Frequencies of Vβ14^+^ and Vβ10b^+^ CD4 T cells in non-proliferating (Ki-67^−^) vs. proliferating (Ki-67^+^) cells from the lungs of flu-infected mice treated with naive or convalescent serum at specified times after infection. (e) UMAP visualization of pooled CD4^+^ T cells from early vs. late convalescent serum treated mice colored by Vβ14 expression. Results from n=7 for PBS, n=14 for naïve serum-treated infected mice, n=7 for convalescent serum-treated infected mice at 4 hours, n=7 for convalescent serum-treated infected mice at 4 days, *P<0.05, **P<0.01, ***P<0.001 by one-way ANOVA, mean ± s.e.m.

**Extended Data Figure 12 | Vβ9+ Treg clonal population does not expand after early convalescent serum therapy.** (a) UMAP of Tregs based on the expression of non-TCR proteins. (b) Heatmap of non-TCR protein expression annotated by cluster and fraction of cells falling into each cluster. (c) UMAP visualization of Tregs colored by the expression Ki-67 from all cohorts of mice. (d) Frequencies of Vβ9^+^ Tregs in non-proliferating (Ki-67^−^) vs. proliferating (Ki-67^+^) cells from the lungs of flu-infected mice treated with naive or convalescent serum at specified times after infection. (e) UMAP visualization of pooled Tregs from early vs. late convalescent serum treated mice colored by Vβ9 expression. Results from n=7 for PBS, n=14 for naïve serum-treated infected mice, n=7 for convalescent serum-treated infected mice at 4 hours, n=7 for convalescent serum-treated infected mice at 4 days, *P<0.05, **P<0.01, ***P<0.001 by one-way ANOVA, mean ± s.e.m.

**Extended Data Figure 13 | Semi-supervised clonotyping assignment script efficiently streamlines high-dimensional single-cell proteomic analysis.** (a) Representative dot plot visualizing T cell classification by variable chain expression by the semi-automated assignment script, colored by Vβ assignment. Blue line = threshold of positivity for each chain. (b) The same visualization as in (a) but colored by Vα assignment. (c) Stacked bar plot of the composition of CD8 T cells from each experiment in this study, colored by clonotype assignment. (d) Left: Stacked bar plot of CD8 T cells from each experiment in this study, colored by Vβ assignment. Right: Stacked bar plot of CD8 T cells with an ‘Unassigned’ Vβ chain, colored by Vα assignment. (e) Left: Stacked bar plot of CD8 T cells from each experiment in this study, colored by Vα assignment. Right: Stacked bar plot of CD8 T cells with an ‘‘Unassigned’ Vα chain, colored by Vβ assignment. (f) Same as (c) for CD4 T cells. (g) Same as (d) for CD4 T cells. (h) Same as (e) for CD4 T cells.

