## Supplemental Material for "A mass cytometry approach to track the evolution of T cell responses during infection and immunotherapy by paired T cell receptor repertoire and T cell differentiation state analysis"

**Extended Data Figure 1.** Expansion of *LADD*-specific CD8 T cells.

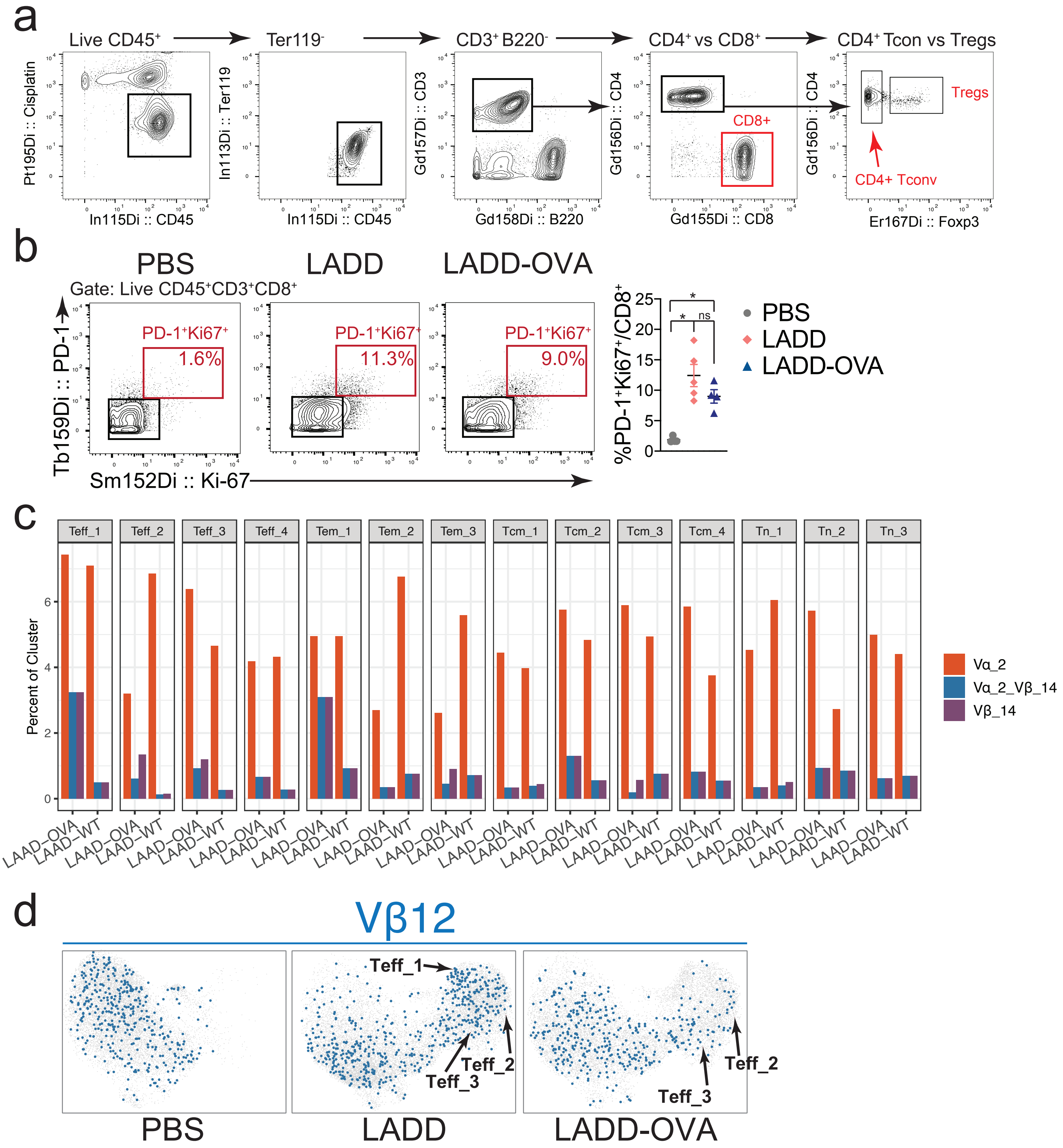

**Extended Data Figure 1 | Tracking a *Listeria*-specific CD8$^+$ T cell response against *LADD* by TCR use and phenotyping with CyTOF.** (a) Gating strategy for all CyTOF analyses preceding input into semi-supervised clonotype assignment script. (b) Representative flow plots and quantification of proliferating CD8$^+$ T cells from spleens 5 days after *LADD, LADD-OVA, or PBS* injection IV. (c) Bar plot of Vα_2, Vβ_14, and Vα_2_Vβ_14 percentages in each cluster for conditions *LAAD* and *LAAD-OVA*. (d) UMAP visualization of pooled CD8$^+$ T cells colored by the expression of Vβ12$^+$ in *LADD*, *LADD-OVA*, or PBS infected mice. For all plots, **P<0.05, **P<0.01, ***P<0.001 by unpaired two-tailed student's T-test, mean ± s.e.m.

**Extended Figure 2.** Expansion and phenotyping of *LADD*-specific CD4 T cell clones from *Listeria*.

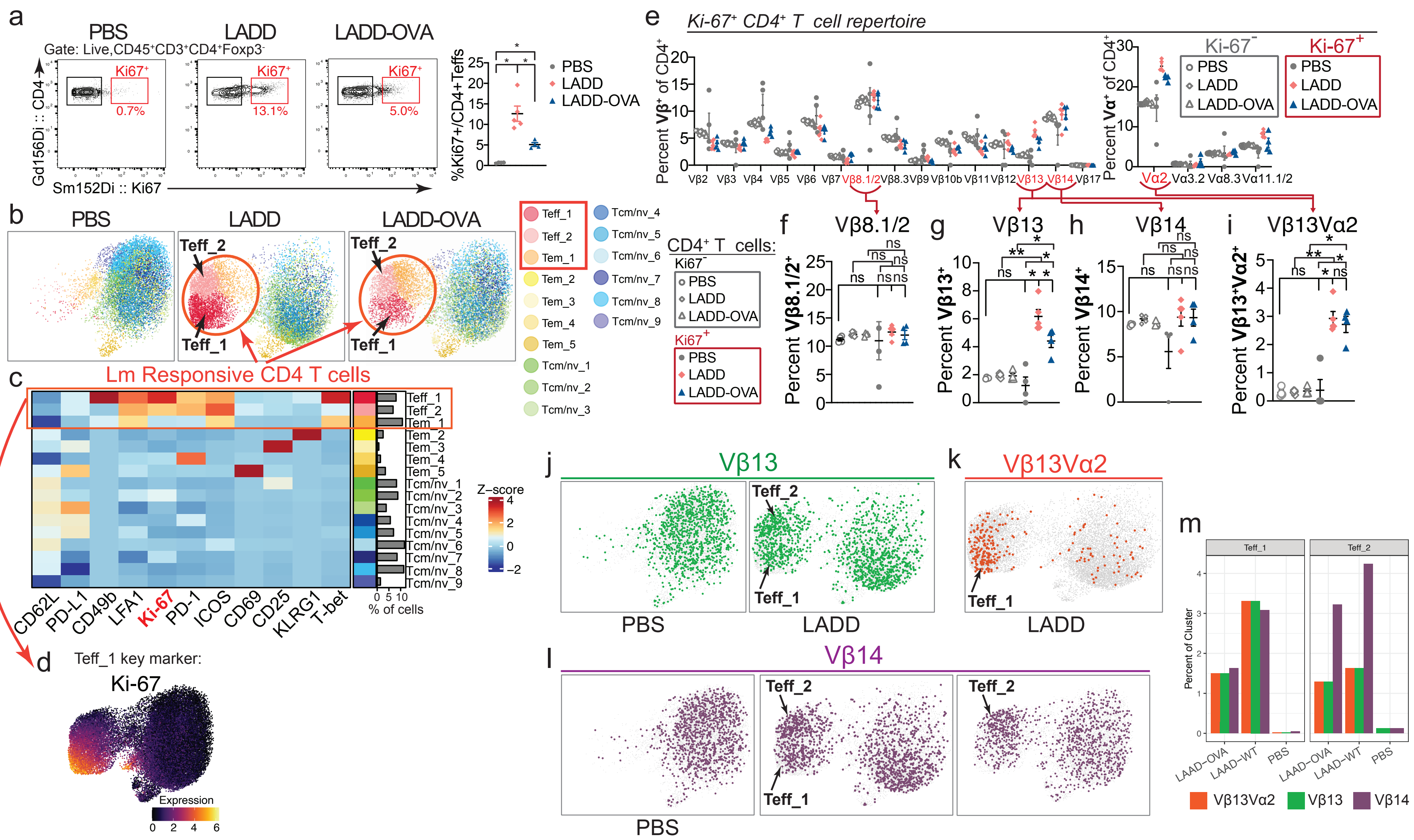

**Extended Data Figure 2 | Tracking *Listeria*-specific CD4$^+$ T cells against *LADD* by TCR use and phenotyping with CyTOF.** (a) Representative flow plots and quantification of proliferating (Ki-67$^+$) CD4$^+$ T cells from spleen 5 days after *LADD, LADD-OVA, or PBS* injection. (b) UMAP visualization of CD4$^+$ T conventional (non-Treg) cell clusters based on expression of non-TCR proteins. (c) Heatmap of non-TCR protein expression annotated by cluster and fraction of cells falling into each cluster. (d) UMAP visualization of CD4$^+$ T cells colored by the expression Ki-67 in *LADD*-infected mice. (e) Frequency of Ki-67$^+$ and Ki-67$^-$ CD4$^+$ T cells using specific TCR Vβ (left) or Vα chains (right) in response to PBS, *LADD*, or *LADD-OVA*. (f-h) Frequency of Vβ13$^+$ (f), Vβ14$^+$ (g), or Vβ13$^+$Vα2$^+$ among Ki-67$^+$(h) versus Ki-67$^-$ CD4$^+$ T cells in *LADD* or *LADD-OVA* infected mice. (i-k) UMAP visualization of pooled CD4$^+$ T cells colored by the expression of Vβ13$^+$ (i), Vβ13$^+$Vα2$^+$ (j), and Vβ14 (k) in *LADD*, *LADD-OVA*, or PBS injected mice. (m) Bar plot of Vβ13$^+$, Vβ13$^+$Vα2$^+$, and Vβ14$^+$ percentages in Teff clusters. Results from n=5 for PBS, n=4 for *LADD*, n=4 for *LADD-OVA*, *P<0.05, **P<0.01, ***P<0.001 by unpaired two-tailed Student's t-test, mean ± s.e.m.

**Extended Data Figure 3.** CD8 response to *LADD*- and OVA-derived epitopes.

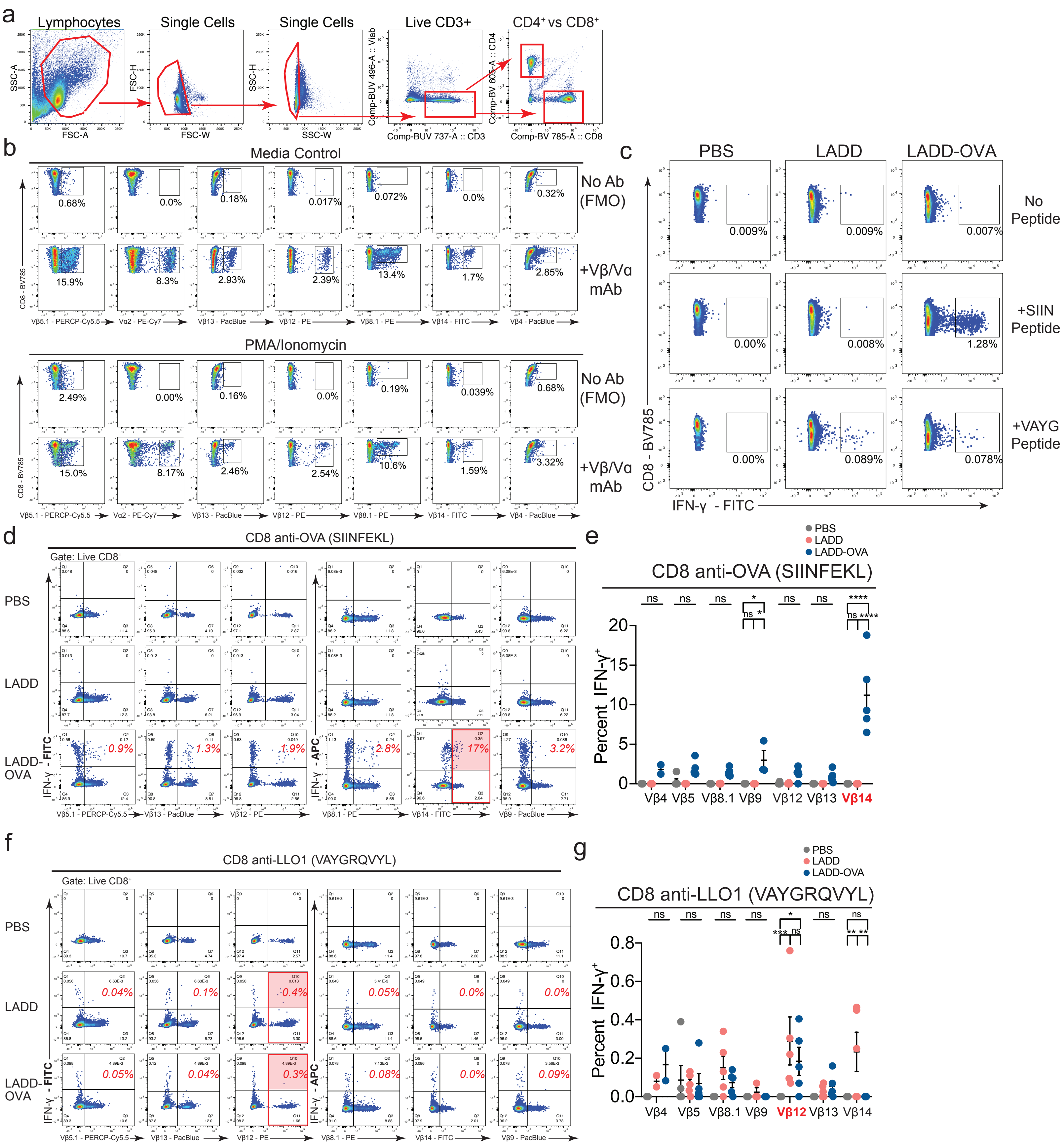

**Extended Data Figure 3 | CD8 response to *LADD*- and OVA-derived epitopes.** (a) Manual gating strategy for all CD8 and CD4 Teff subsets. (b) FMO controls for all fluorescent-conjugated Vβ and Vα antibodies in naive splenocytes +/- stimulation with PMA/Ionomycin. (c) Frequency of IFN-$y^+$ CD8s in splenocytes in unstimulated controls versus peptide conditions. (d) Representative flow plots of IFN-$y^+$ CD8s versus respective Vβ chains in splenocytes stimulated with SIINFEKL peptide. (e) Percent of IFN-$y^+$ for respective Vβ$^+$ CD8s in splenocytes stimulated with SIINFEKL peptide. (f) Representative flow plots of IFN-$y^+$ CD8s versus respective Vβ chains in splenocytes stimulated with LLO1 peptide. (g) Percent of IFN-$y^+$ for respective Vβ$^+$ CD8 in splenocytes stimulated with LLO1 peptide. Results from n=5 for PBS, n=4 for *LADD*, n=4 for *LADD-OVA*, *P<0.05, **P<0.01, ***P<0.001 by unpaired two-tailed Student's t-test, mean ± s.e.m.

### **Extended Data Figure 4.** CD4 response to *LADD*-derived antigens.

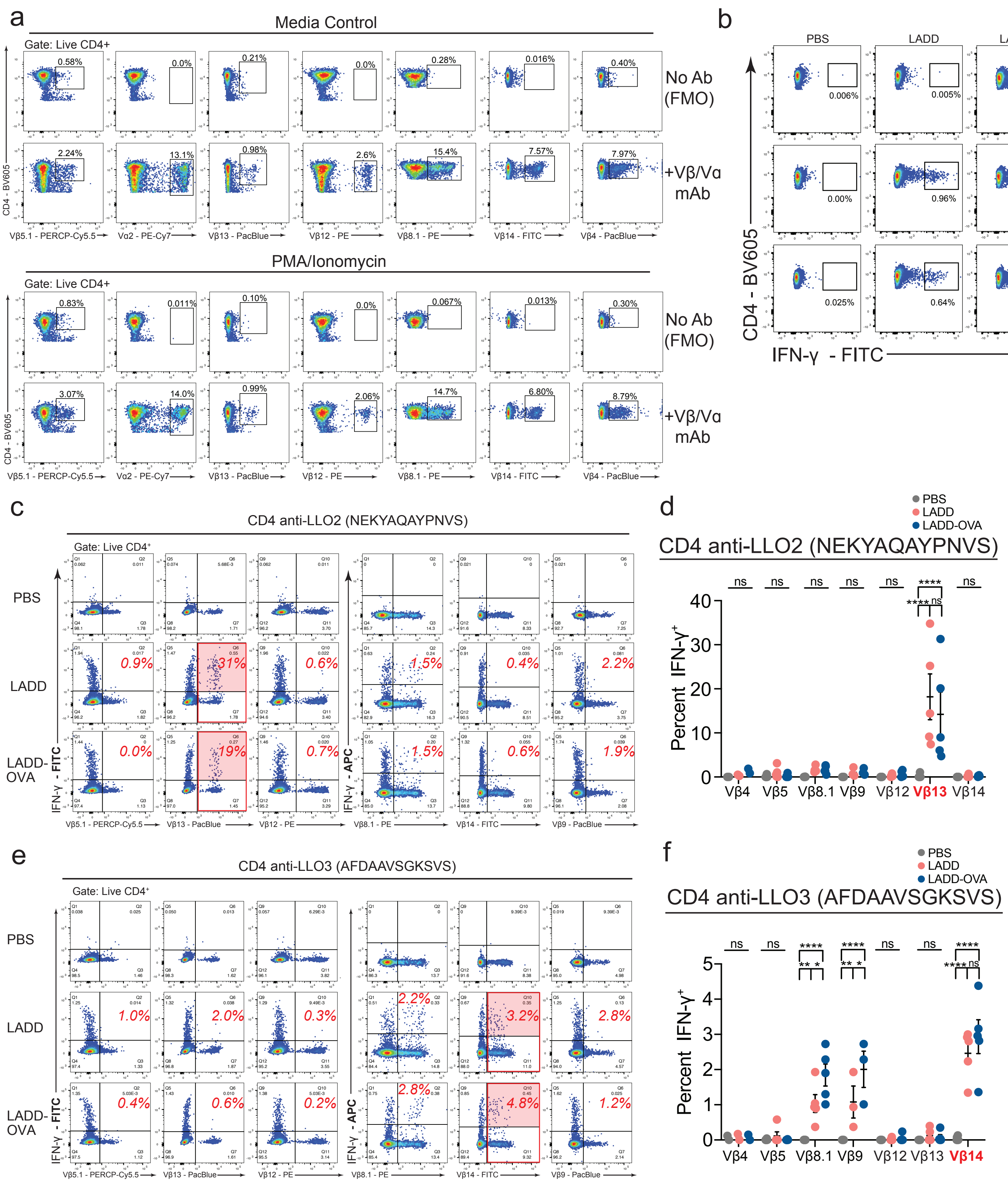

**Extended Data Figure 4 | CD4 response to *LADD*-derived epitopes.** (a) FMO controls for all fluorescent-conjugated Vβ and Vα antibodies in naive splenocytes +/- stimulation with PMA/Ionomycin. (b) Frequency of IFN-y$^+$ CD4s in splenocytes in unstimulated controls versus peptide conditions. (c) Representative flow plots of IFN-y$^+$ CD4s versus respective Vβ chains in splenocytes stimulated with LLO2 peptide. (d) Percent of IFN-y$^+$ for respective Vβ$^+$ CD4s in splenocytes stimulated with LLO2 peptide. (e) Representative flow plots of IFN-y$^+$ CD4s versus respective Vβ chains in splenocytes stimulated with LLO3 peptide. (f) Percent of IFN-y$^+$ for respective Vβ$^+$ CD4s in splenocytes stimulated with LLO3 peptide. Results from n=5 for PBS, n=4 for *LADD*, n=4 for *LADD-OVA*, *P<0.05, **P<0.01, ***P<0.001 by unpaired two-tailed Student's t-test, mean ± s.e.m.

**Extended Data Figure 5.** Validation of Vβ14$^{+}$Vα2$^{+}$ CD8 expansion with scRNA-seq.

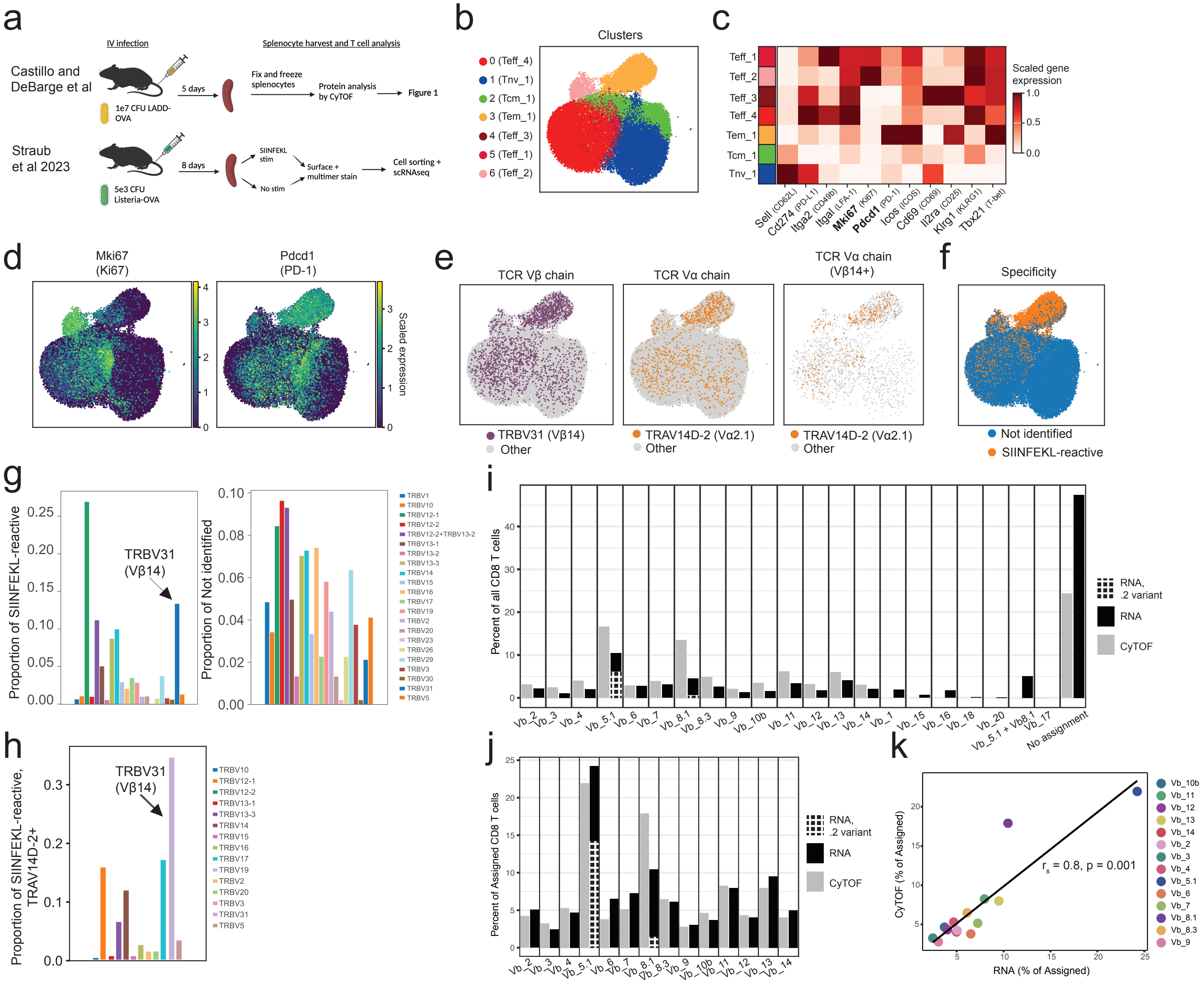

**Extended Data Figure 5 | Comparative analysis of Vβ14+Vα2+ CD8 expansion with scRNAseq during *Listeria* infection.** *(a)* Schematic comparison of approach used in this study compared to Straub et al 2023. (b) Re-annotated clusters identified by Straub et al plotted in UMAP space. (c) Scaled gene expression within the scRNA-seq dataset of genes encoding the proteins measured in Fig. 1, stratified by cluster. (d) UMAP visualization of all clustered cells colored by Ki67 or PD-1 gene expression. (e) UMAP visualization of all clustered cells colored by TRBV31 assignment or TRAV14D-2 assignment (**left**), and UMAP visualization of TRBV31 assigned cells colored by TRAV14D-2 assignment (**right**). (f) UMAP visualization of all clustered cells colored by identified specificity. (g) Quantification of TRBV gene usage among SIINFEKL-reactive cells (right) or of cells of unidentified reactivity (left). (h) Quantification of TRBV gene usage among SIINFEKL- reactive cells expressing TRAV14D-2. (i) Quantification of TRBV/Vβ chain assignment of all CD8 T cells as measured by CyTOF or scRNA-seq. (j) Quantification of the TRBV/Vβ chains measured in the CyTOF assay of all CD8 T cells that were assigned a TRBV/Vβ chain by either CyTOF or scRNA-seq. (k) Spearman correlation analysis of data shown in (j).

**Extended Data Figure 6.** Activation and proliferation of CD8, CD4, and Treg cells in response to flu across tissues.

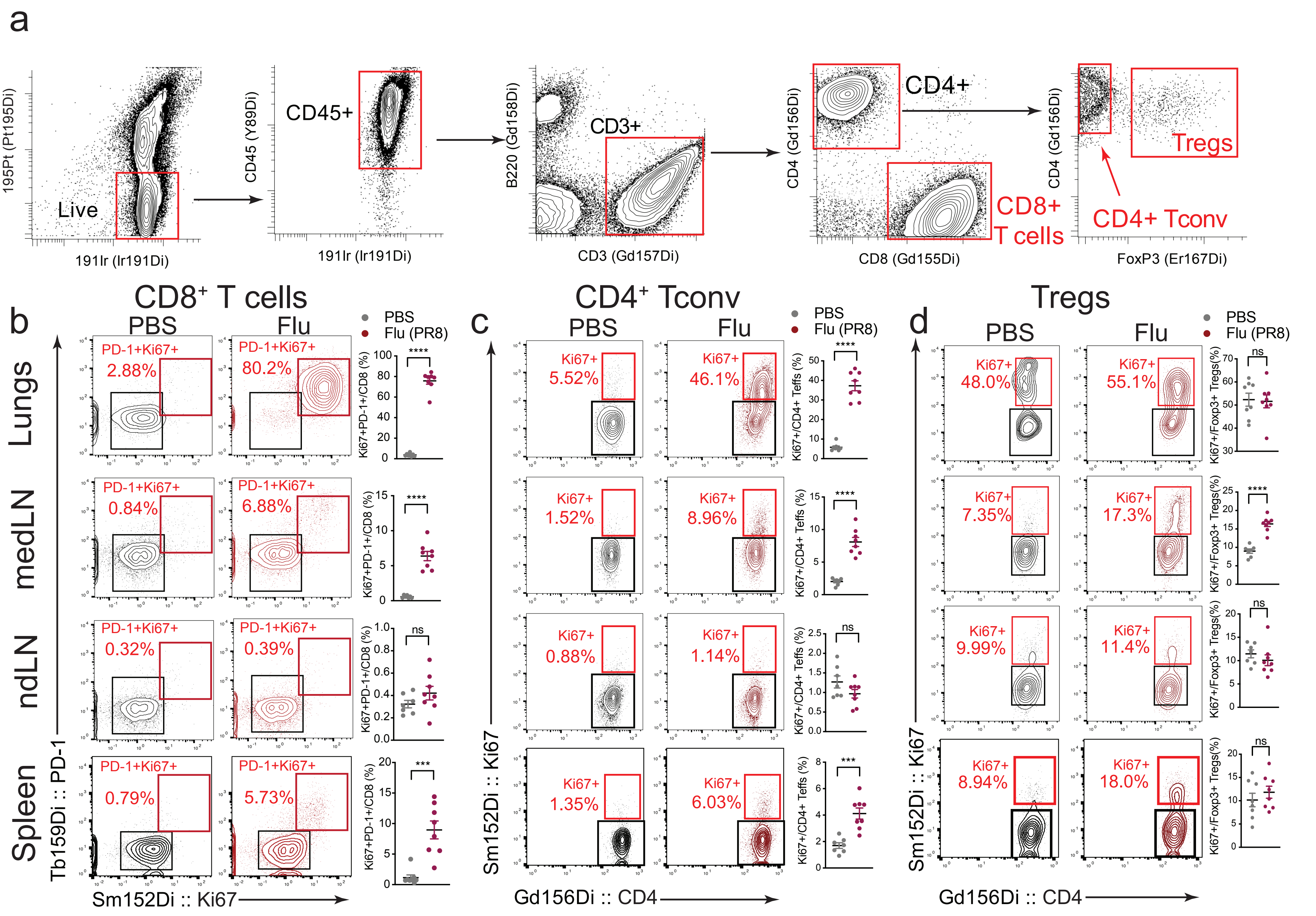

**Extended Data Figure 6 | Expansion and activation of T cell subsets during flu infection across multiple tissues.** (a) Gating strategy for all CyTOF analyses preceding input into semi-supervised clonotype assignment script. (b) Representative flow plots and quantification of proliferating (PD-$1^{+}$Ki$67^{+}$) $CD8^{+}$ T cells. (c) Representative flow plots and quantification of proliferating (Ki$67^{+}$) Foxp$3^{-}$ $CD4^{+}$ Teff cells. (d) Representative flow plots and quantification of proliferating (Ki$67^{+}$) Foxp$3^{+}$ Tregs. For all plots, **$P<0.05$, **$P<0.01$, ***$P<0.001$ by one-way ANOVA, mean ± s.e.m.

**Extended Data Figure 7.** Tracking of CD4 clonal expansion and recognition of flu antigens.

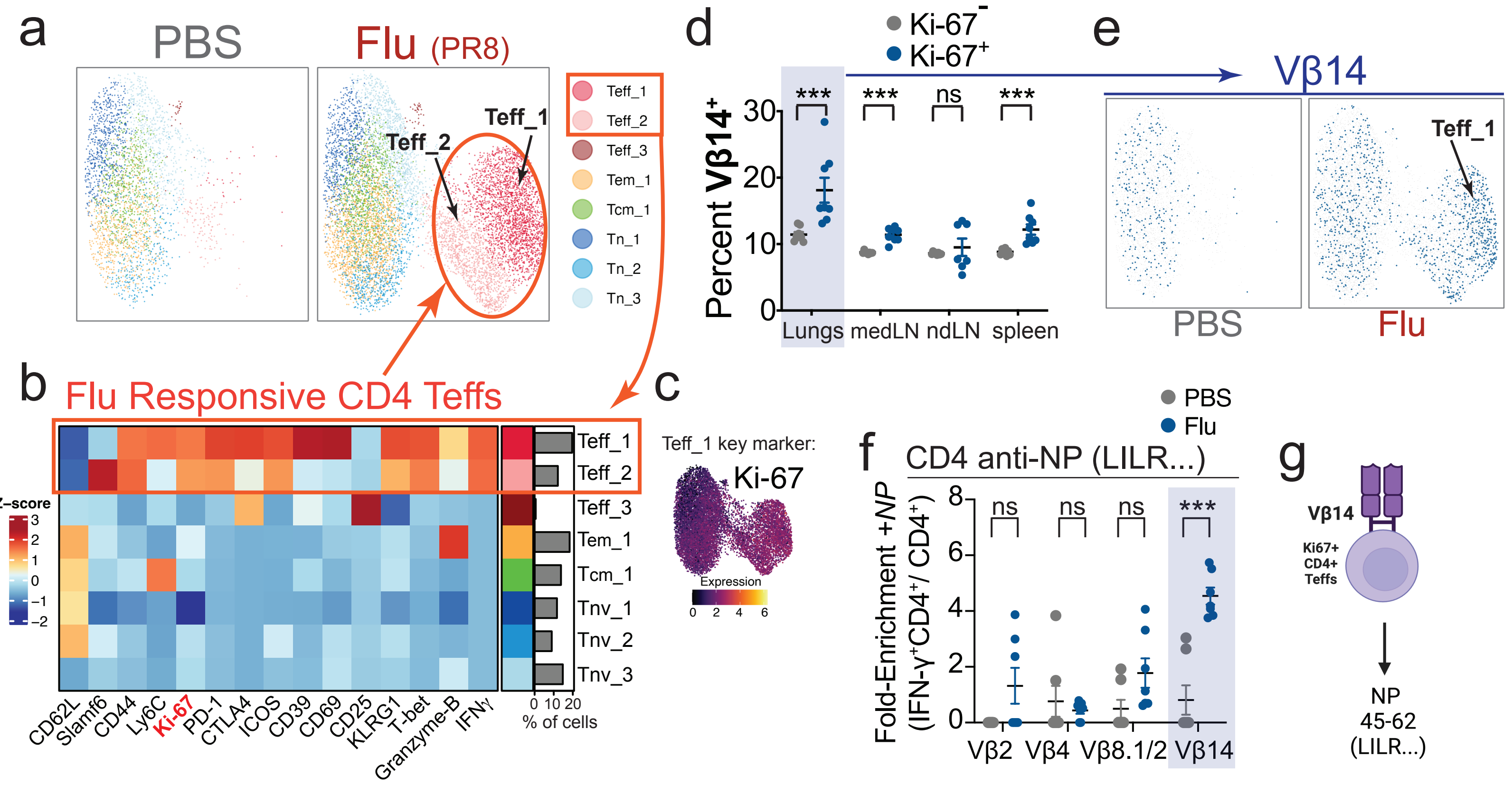

**Extended Data Figure 7 | Identification of flu NP-specific Vβ14$^+$ CD4$^+$ T cells.** (a) UMAP visualization of CD4$^+$ T conventional (non-Treg) cell clusters based on expression of non-TCR proteins. (b) Heatmap of non-TCR protein expression annotated by cluster and fraction of cells falling into each cluster. (c) UMAP visualization of CD4$^+$ T cells colored by the expression Ki-67 in flu-infected mice. (d) Frequencies of Vβ14$^+$ CD4 T cells in non-proliferating (Ki-67$^-$) vs. proliferating (Ki-67$^+$) cells across indicated tissues. (e) UMAP visualization of pooled CD4$^+$ T cells colored by Vβ14 expression. (f) Quantification of fold-enrichment in Vβ-chain usage by CD4$^+$ T cells stimulated with the PR8 NP peptide (LILRGSVAHKSCLPACV). (g) Schematic summarizing that Vβ14$^+$ CD4$^+$ T cells are enriched to recognize PR8 NP antigen from flu. Results from n=7 for PBS, n=7 for PR8, *$P<0.05$, **$P<0.01$, ***$P<0.001$ by unpaired two-tailed Student's t-test, mean ± s.e.m.

**Extended Data Figure 8**. Expansion of eTregs after influenza infection.

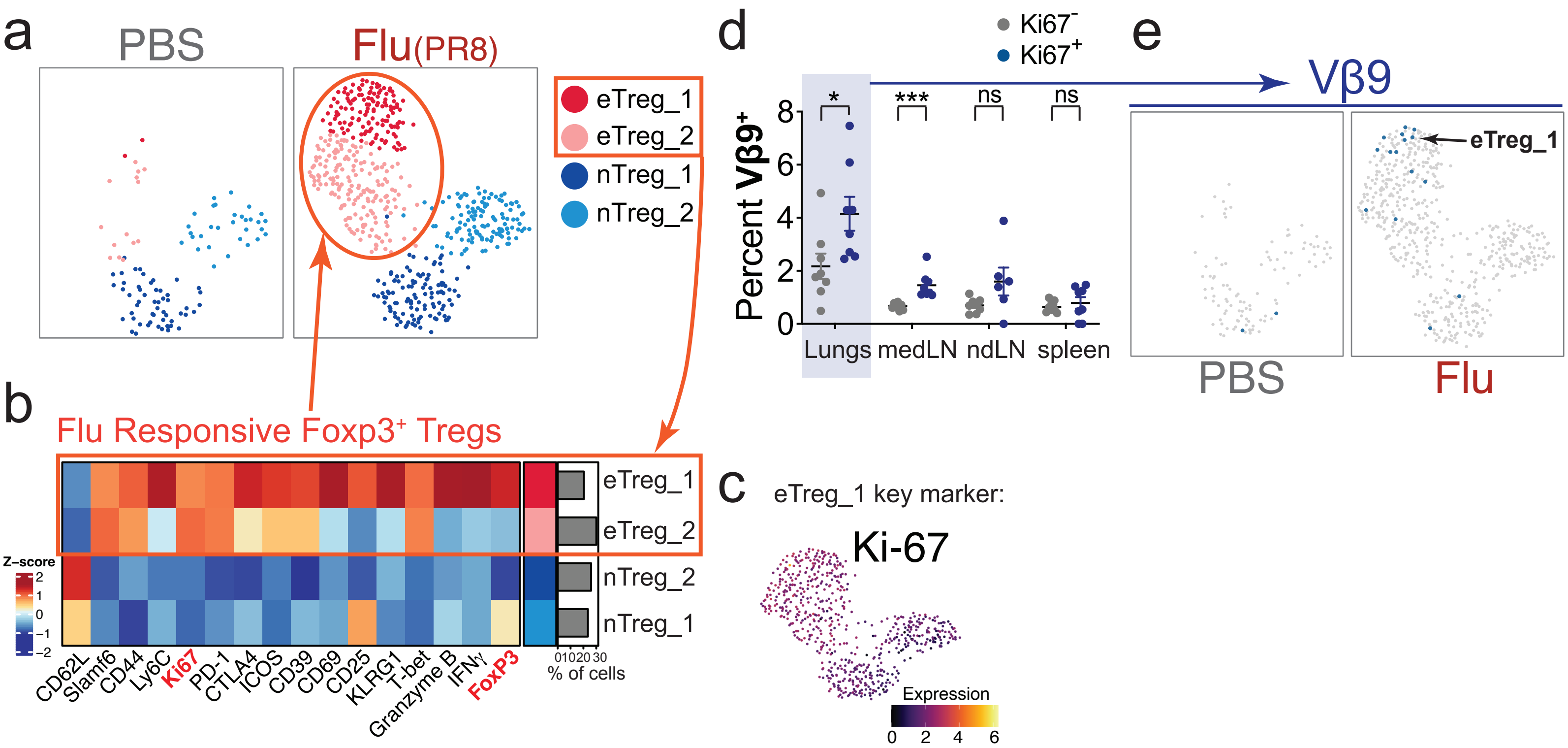

**Extended Data Figure 8 | Identification and phenotyping of Vβ9+ Treg expansion after flu infection.** (a) UMAP of Treg populations based on expression of non-TCR proteins. (b) Heatmap of non-TCR protein expression annotated by cluster and fraction of cells falling into each cluster. (c) UMAP visualization of Tregs colored by the expression Ki-67 in flu-infected mice. (d) Frequencies of Vβ9$^+$ Tregs in non-proliferating (Ki-67$^-$) vs. proliferating (Ki-67$^+$) cells across indicated tissues. (e) UMAP visualization of pooled Tregs from PBS or flu-infected mice colored by Vβ9 expression. Results from n=7 for PBS, n=7 for PR8, *P<0.05, **P<0.01, ***P<0.001 by unpaired two-tailed Student's t-test, mean ± s.e.m.

**Extended Data Figure 9.** Tracking CD4 T cell clonal expansion during flu vaccination.

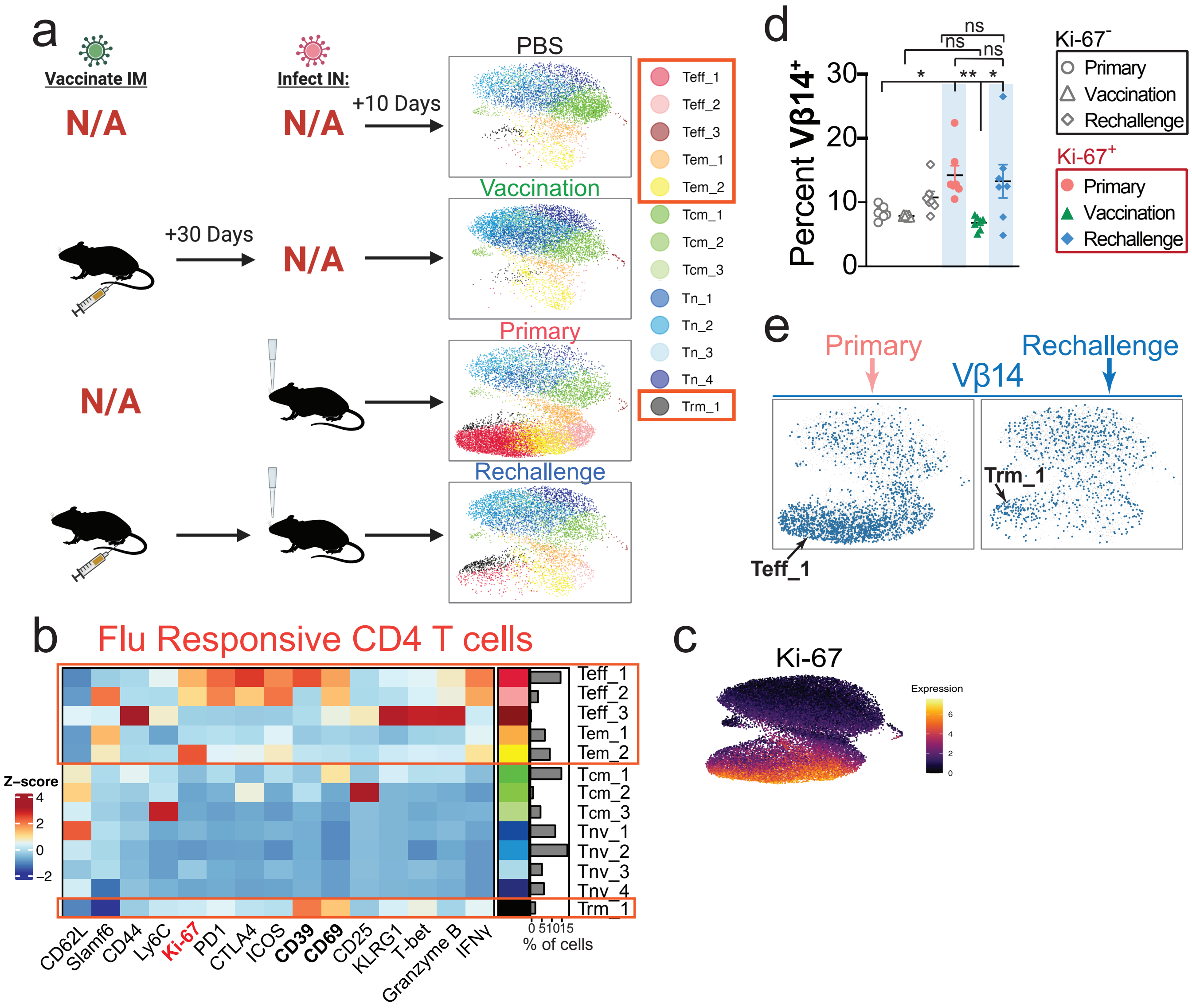

**Extended Data Figure 9 | *Flu*-specific conventional CD4$^+$ T cell differentiation, but not repertoire, is altered by vaccination.** (a) UMAP visualization of CD4$^+$ T conventional (non-Treg) cell clusters based on expression of non-TCR proteins. (b) Heatmap of non-TCR protein expression annotated by cluster and fraction of cells falling into each cluster. (c) UMAP visualization of CD4$^+$ T cells colored by the expression Ki-67 from all cohorts of mice. (d) Frequencies of Vβ14$^+$ CD4 T cells in non-proliferating (Ki-67$^-$) vs. proliferating (Ki-67$^+$) cells in lungs of primary, vaccinated, or rechallenged mice. (e) UMAP visualization of pooled CD4$^+$ T cells from primary or rechallenged mice colored by Vβ14 expression. Results from n=7 for PBS, n=7 for PR8, *P<0.05, **P<0.01, ***P<0.001 by one-way ANOVA, mean ± s.e.m.

**Extended Data Figure 10.** Tracking Treg clonal expansion during flu vaccination.

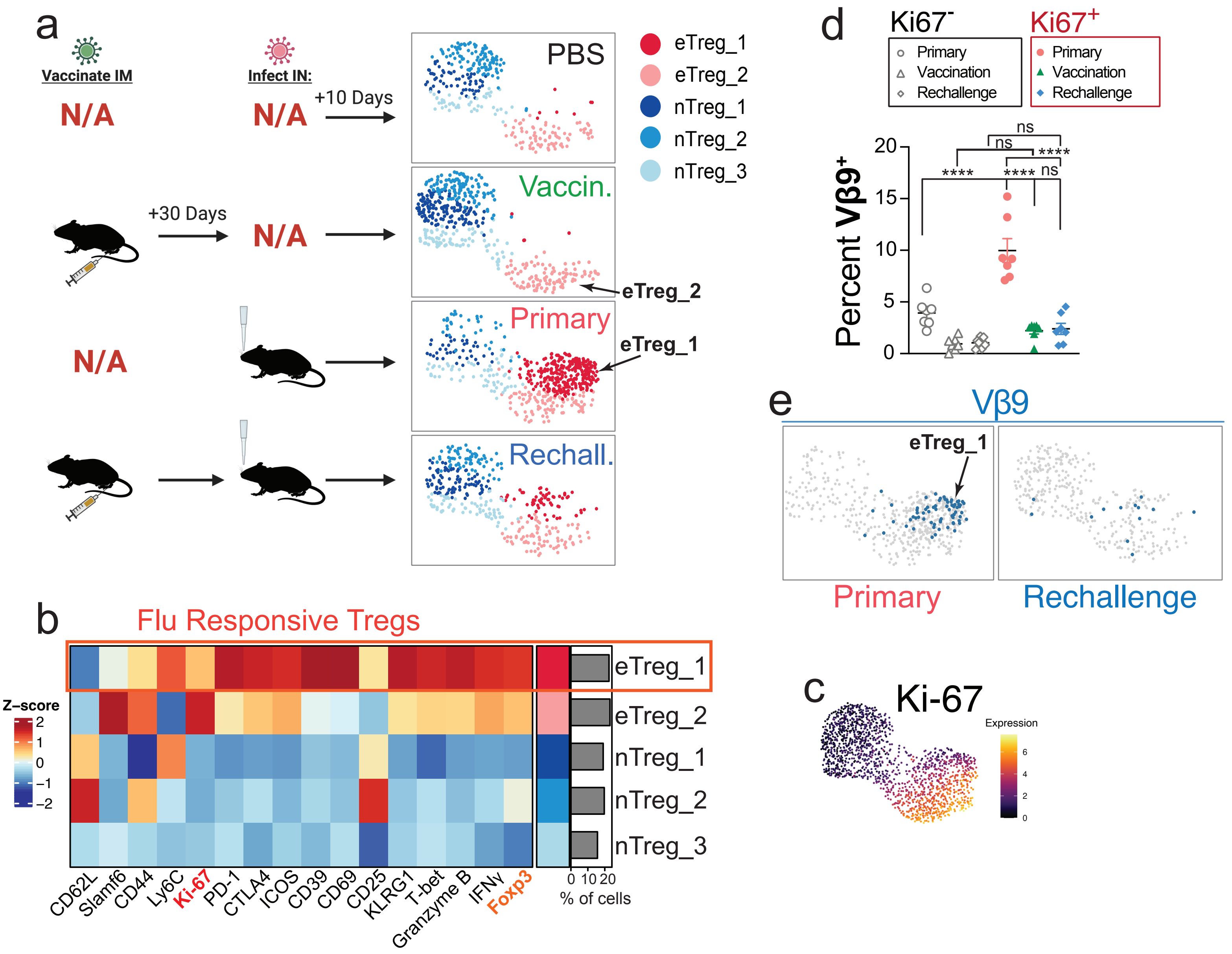

**Extended Figure 10 | Vβ9+ Treg clonal population is only present in the setting of a primary infection.** (a) UMAP of Treg populations based on the expression of non-TCR proteins. (b) Heatmap of non-TCR protein expression annotated by cluster and fraction of cells falling into each cluster. (c) UMAP visualization of Tregs colored by the expression Ki-67 from all cohorts of mice. (d) Frequencies of $V\beta9^{+}$ Tregs in non-proliferating ($Ki\text{-}67^{-}$) vs. proliferating ($Ki\text{-}67^{+}$) cells in lungs of primary, vaccinated, or rechallenged mice. (e) UMAP visualization of pooled Tregs from primary or rechallenged mice colored by Vβ9 expression. Results from n=7 for PBS, n=7 for PR8, *$P<0.05$, **$P<0.01$, ***$P<0.001$ by one-way ANOVA, mean ± s.e.m.

**Extended Data Figure 11.** Minimal impact on CD4 T cell response with addition of serum.

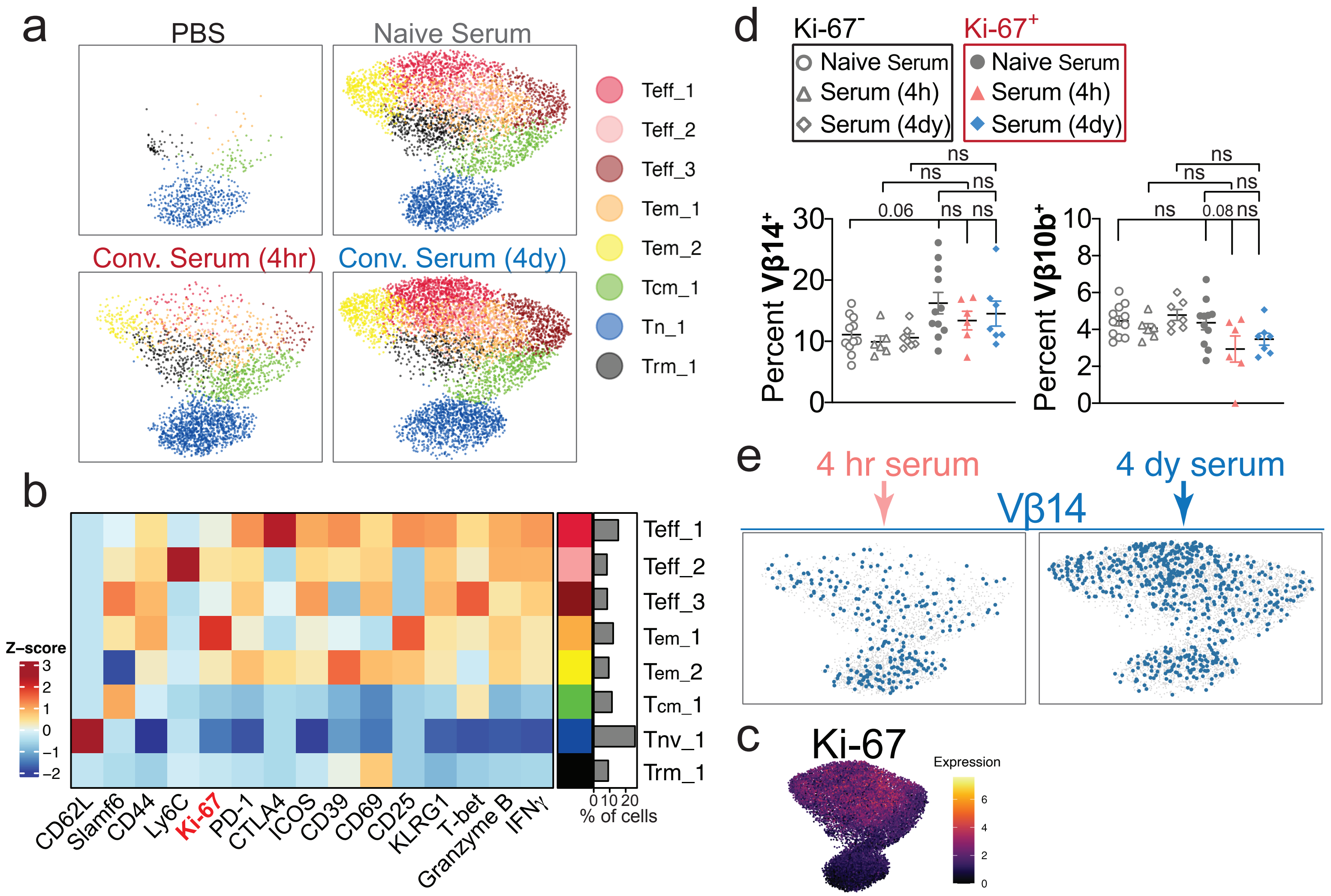

**Extended Data Figure 11 | Convalescent therapy has a limited effect on the CD4$^+$ Teff repertoire during influenza infection.** (a) UMAP visualization of CD4$^+$ T conventional (non-Treg) cell clusters based on expression of non-TCR proteins. (b) Heatmap of non-TCR protein expression annotated by cluster and fraction of cells falling into each cluster. (c) UMAP visualization of CD4$^+$ T cells colored by the expression Ki-67 from all cohorts of mice. (d) Frequencies of Vβ14$^+$ and Vβ10b$^+$ CD4 T cells in non-proliferating (Ki-67$^-$) vs. proliferating (Ki-67$^+$) cells from the lungs of flu-infected mice treated with naive or convalescent serum at specified times after infection. (e) UMAP visualization of pooled CD4$^+$ T cells from early vs. late convalescent serum treated mice colored by Vβ14 expression. Results from n=7 for PBS, n=14 for naïve serum-treated infected mice, n=7 for convalescent serum-treated infected mice at 4 hours, n=7 for convalescent serum-treated infected mice at 4 days, *$P<0.05$, **$P<0.01$, ***$P<0.001$ by one-way ANOVA, mean ± s.e.m.

**Extended Data Figure 12.** Tracking Treg clonal expansion against flu after with serum transfer.

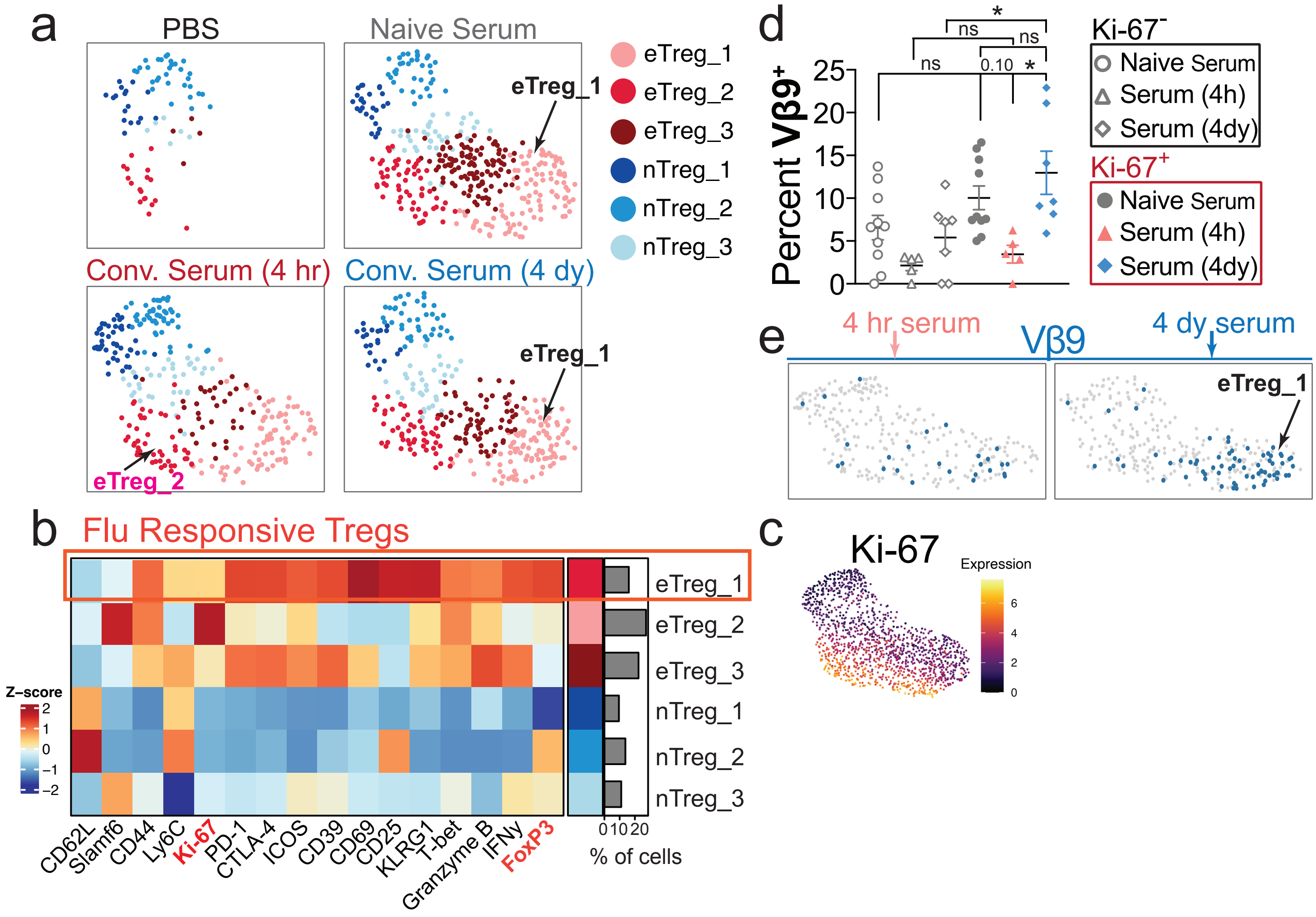

**Extended Data Figure 12 | Vβ9+ Treg clonal population does not expand after early convalescent serum therapy.** (a) UMAP of Tregs based on the expression of non-TCR proteins. (b) Heatmap of non-TCR protein expression annotated by cluster and fraction of cells falling into each cluster. (c) UMAP visualization of Tregs colored by the expression Ki-67 from all cohorts of mice. (d) Frequencies of Vβ9$^+$ Tregs in non-proliferating (Ki-67$^-$) vs. proliferating (Ki-67$^+$) cells from the lungs of flu-infected mice treated with naive or convalescent serum at specified times after infection. (e) UMAP visualization of pooled Tregs from early vs. late convalescent serum treated mice colored by Vβ9 expression. Results from n=7 for PBS, n=14 for naïve serum-treated infected mice, n=7 for convalescent serum-treated infected mice at 4 hours, n=7 for convalescent serum-treated infected mice at 4 days, *$P<0.05$, **$P<0.01$, ***$P<0.001$ by one-way ANOVA, mean ± s.e.m.

**Extended Data Figure 13.** Semi-supervised clonotyping assignment of high dimensional single-cell proteomic data.

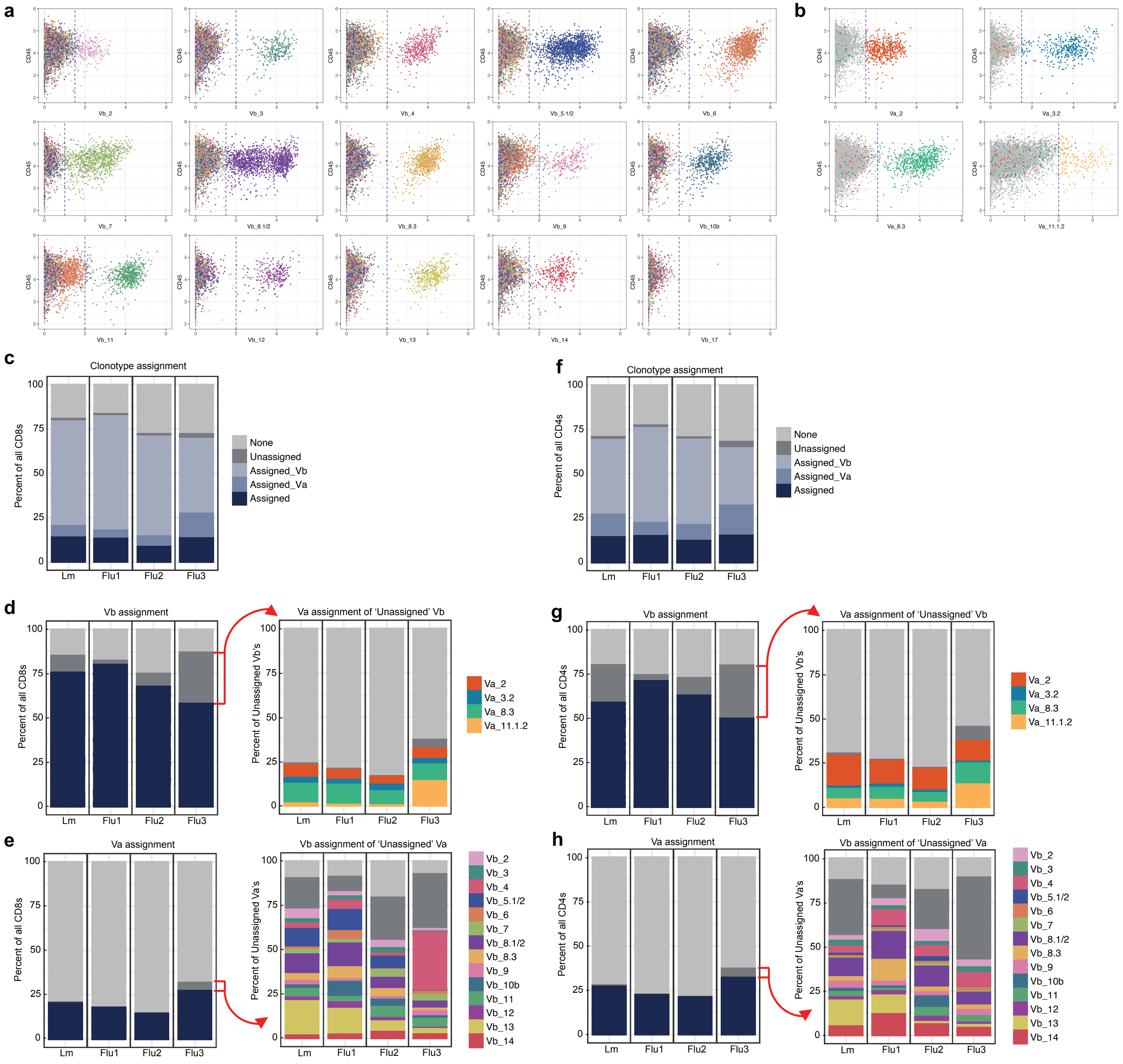

**Extended Data Figure 13 | Semi-supervised clonotyping assignment script efficiently streamlines high-dimensional single-cell proteomic analysis.** (a) Representative dot plot visualizing T cell classification by variable chain expression by the semi-automated assignment script, colored by Vβ assignment. Blue line = threshold of positivity for each chain. (b) The same visualization as in (a) but colored by Vα assignment. (c) Stacked bar plot of the composition of CD8 T cells from each experiment in this study, colored by clonotype assignment. (d) Left: Stacked bar plot of CD8 T cells from each experiment in this study, colored by Vβ assignment. Right: Stacked bar plot of CD8 T cells with an 'Unassigned' Vβ chain, colored by Vα assignment. (e) Left: Stacked bar plot of CD8 T cells from each experiment in this study, colored by Vα assignment. Right: Stacked bar plot of CD8 T cells with an ''Unassigned' Vα chain, colored by Vβ assignment. (f) Same as (c) for CD4 T cells. (g) Same as (d) for CD4 T cells. (h) Same as (e) for CD4 T cells.

| Channel | Marker | Vendor | Cat Number | Clone | Species | [Optimal] (ug/mL) | Corresponding figures |
|---|---|---|---|---|---|---|---|
| 89Y | CD45 | Biolegend | 103102 | 30-F11 | Mouse | 6 | Fig 4, 5, 6 |
| 113In | Ter119 | Biolegend | 116202 | TER119 | Mouse | 3 | Fig 1, 4, 5, 6 |
| 115In | CD45 | Biolegend | 103102 | 30-F11 | Mouse | 3 | Fig 1 |
| 139La | CD44 | Biolegend | 103002 | IM7 | Mouse | 0.75 | Fig 4, 5, 6 |
| 139La | Ly6G | Biolegend | 127602 | 1A8 | Mouse | 1.5 | Fig 1 |
| 140Ce | KLRG1 | R&D Systems | MAB69441-100 | 2F1 | Mouse | 3 | Fig 1, 4, 5, 6 |
| 141Pr | Granzyme B | Biolegend | 372202 | QA16A02 | Mouse | 3 | Fig 4, 5, 6 |
| 141Pr | CD11b | Biolegend | 101202 | M1/70 | Mouse | 1.5 | Fig 1 |
| 142Nd | Ly6C | Biolegend | 128002 | HK1.4 | Mouse | 0.75 | Fig 4, 5, 6 |
| 142Nd | CD49b | Biolegend | 103501 | HMa2 | Mouse | 0.1875 | Fig 1 |
| 143Nd | Vb10b | BD | Custom order | B21.5 | Mouse | 0.75 | Fig 1, 4, 5, 6 |
| 144Nd | Va2 | BD | Custom order | B20.1 | Mouse | 0.75 | Fig 1, 4, 5, 6 |
| 145Nd | CD39 | Thermo Scientific | 14-0391-82 | 24DMS1 | Mouse | 0.75 | Fig 4, 5, 6 |
| 145Nd | NK1.1 | Biolegend | 108702 | PK136 | Mouse | 3 | Fig 1 |
| 146Nd | Va11.1/11.2 | BD | Custom order | RR8-1 | Mouse | 6 | Fig 1, 4, 5, 6 |
| 147Sm | IFNy | Biolegend | 505834 | XMG1.2 | Mouse | 3 | Fig 4, 5, 6 |
| 147Sm | PD-L1 | Biolegend | 135202 | 10F.9G2 | Mouse | 3 | Fig 1 |
| 148Nd | Vb14 | BD | Custom order | 14-2 | Mouse | 3 | Fig 1, 4, 5, 6 |
| 149Sm | Vb8.3 | BD | Custom order | 1B3.3 | Mouse | 3 | Fig 1, 4, 5, 6 |
| 150Nd | Vb4 | BD | Custom order | KT4 | Mouse | 3 | Fig 1, 4, 5, 6 |
| 151Eu | Vb12 | BD | Custom order | MR11-1 | Mouse | 0.75 | Fig 1, 4, 5, 6 |
| 152Sm | Ki67 | Thermo Scientific | 12-5698-82 | SolA15 | Mouse | 6 | Fig 1, 4, 5, 6 |
| 153Eu | Vb8.1/2 | BD | Custom order | MR5-2 | Mouse | 3 | Fig 1, 4, 5, 6 |
| 154Sm | Vb5.1/2 | BD | Custom order | MR9-4 | Mouse | 3 | Fig 1, 4, 5, 6 |
| 155Gd | CD8 | Biolegend | 100702 | 53-6.7 | Mouse | 3 | Fig 1, 4, 5, 6 |
| 156Gd | CD4 | Biolegend | 100506 | RM4-5 | Mouse | 0.75 | Fig 1, 4, 5, 6 |
| 157Gd | CD3 | Biolegend | 100202 | 17A2 | Mouse | 0.75 | Fig 1, 4, 5, 6 |
| 158Gd | B220 | Biolegend | 103202 | RA3-6B2 | Mouse | 1.5 | Fig 1, 4, 5, 6 |
| 159Tb | PD-1 | Biolegend | 135202 | 29.F.1A12 | Mouse | 0.75 | Fig 1, 4, 5, 6 |
| 160Gd | Vb3 | BD | Custom order | KJ25 | Mouse | 0.375 | Fig 1, 4, 5, 6 |
| 161Dy | Tbet | Biolegend | 644802 | 4B10 | Mouse | 6 | Fig 1, 4, 5, 6 |
| 162Dy | Ly108 (Slamf6) | Thermo Scientific | 14-1508-82 | 13G3-19D | Mouse | 1.5 | Fig 4, 5, 6 |
| 162Dy | TCRgd | Biolegend | 118101 | GL3 | Mouse | 3 | Fig 1 |
| 163Dy | Va8.3 | BD | Custom order | B21.14 | Mouse | 3 | Fig 1, 4, 5, 6 |
| 164Dy | Vb2 | BD | Custom order | B20.6 | Mouse | 3 | Fig 1, 4, 5, 6 |
| 165Ho | CD69 | R&D Systems | AF2386 | polyclonal | Mouse | 0.75 | Fig 1, 4, 5, 6 |
| 166Er | Vb17 | BD | Custom order | KJ23 | Mouse | 0.75 | Fig 1, 4, 5, 6 |
| 167Er | FoxP3 | Thermo Scientific | 14-4771-80 | NRRF-30 | Mouse | 3 | Fig 1, 4, 5, 6 |
| 168Er | CD25 | Biolegend | 102040 | PC61 | Mouse | 3 | Fig 1, 4, 5, 6 |
| 169Tm | CTLA-4 | Biolegend | 106302 | UC10-4B9 | Mouse | 6 | Fig 4, 5, 6 |
| 169Tm | CD62L-FITC | Biolegend | 408302 (a-FITC) | CD62L: MEL14 | Mouse | then 6ug/mL a- | Fig 1 |
| 170Er | Vb7 | BD | Custom order | TR310 | Mouse | 0.75 | Fig 1, 4, 5, 6 |
| 171Yb | ICOS | Biolegend | 313502 | C398.4A | Mouse | 3 | Fig 1, 4, 5, 6 |
| 172Yb | Vb9 | BD | Custom order | MR10-2 | Mouse | 0.375 | Fig 1, 4, 5, 6 |
| 173Yb | Vb6 | BD | Custom order | RR4-7 | Mouse | 1.5 | Fig 1, 4, 5, 6 |
| 174Yb | Vb11 | BD | Custom order | RR3-15 | Mouse | 0.75 | Fig 1, 4, 5, 6 |
| 175Lu | Vb13 | BD | Custom order | MR12-3 | Mouse | 0.75 | Fig 1, 4, 5, 6 |
| 176Yb | Va3.2 | BD | Custom order | RR3-16 | Mouse | 0.75 | Fig 1, 4, 5, 6 |
| 209Bi | MHCII | Biolegend | 107602 | M5/114.15.2 | Mouse | 0.1875 | Fig 1 |
| 209Bi | CD62L | Fisher Scientific | MAB5761SP | 95218 | Mouse | 6 | Fig 4, 5, 6 |

| **Conventional T cell cluster family** | **Cluster defining markers** | **Variable markers (among clusters of the same family)** |
|---|---|---|
| Naive | CD62L+ CD44- | Slamf6 |
| Central memory | CD62L+ CD44+ | Slamf6, Ki67, ICOS, Ly6C |
| Effector memory | CD62L- CD44+ KLRG1- | Tbet, CD25, ICOS, Ki67, PD-1, IFNy, Granzyme B, Ly6C, Slamf6, CD39, CD69, CTLA4 |
| Effectors | CD62L- CD44- KLRG1+ Tbet+ | LFA-1, CD25, ICOS, Ki67, PD-1, IFNy, Granzyme B, Ly6C, Slamf6, CD39, CD69, CTLA4 |
